# Harnessing intrinsic fluorescence for typing of secondary structures of DNA

**DOI:** 10.1101/2020.01.15.907501

**Authors:** Michela Zuffo, Aurélie Gandolfini, Brahim Heddi, Anton Granzhan

## Abstract

DNA is polymorphic since, despite its ubiquitous presence as a double-stranded helix, it is able to fold into a plethora of other secondary structures both *in vitro* and in cells. Despite the considerable advances that have been made in understanding this structural diversity, its high-throughput investigation still faces severe limitations. This mainly stems from the lack of suitable label-free methods, combining a fast and cheap experimental workflow with high information content. Here, we explore the use of intrinsic fluorescence emitted by nucleic acids for this scope. After a preliminary assessment of the suitability of this phenomenon for tracking the conformational changes of DNA, we examined the intrinsic steady-state emission spectra of an 89-membered set of synthetic oligonucleotides with reported conformation (G-quadruplexes, i-motifs, single- and double stranded DNA) by means of multivariate analysis. Specifically, principal component analysis of emission spectra resulted in successful clustering of oligonucleotides into three corresponding conformational groups, albeit without discrimination between single- and double-stranded structures. Linear discriminant analysis of the same training set was exploited for the assessment of new sequences, allowing the evaluation of their G4-forming propensity. Our method does not require any labelling agent or dye, avoiding the related intrinsic bias, and can be utilized to screen novel sequences of interest in a high-throughput and cost-effective manner. In addition, we observed that left-handed (Z-) G4 structures were systematically more fluorescent than most other G4 structures, almost reaching the quantum yield of 5′-d[(G_3_T)_3_G_3_]-3′ (*G*_*3*_*T*), the most fluorescent G4 structure reported to date. This property is likely to arise from the similar base-stacking geometry in both types of structures.

## INTRODUCTION

Since the elucidation of the double-helical structure of DNA, a great deal of effort has been devoted to understand whether the genomic material could also adopt other conformations. It is now firmly established that DNA can fold into a wealth of secondary structures that are intertwined in delicate equilibria. These range from single-, double- and triple-stranded conformations (i.e., random-coil, A-, B- and Z-DNA, and triplexes, respectively) to three- and four-way junctions as well as tetra-stranded structures (chiefly, G-quadruplexes, or G4, and i-motifs, or iM). Specifically, G4 structures are formed by stacks of guanine quartets, stabilized by monovalent cations (mostly K^+^ and Na^+^). Despite this common scaffold, G4 structures are themselves extremely polymorphic in terms of strand topology (parallel, anti-parallel, hybrid), loop geometry, groove size, etc. (1). On the other hand, iMs consist of two interpenetrated duplexes formed by hemi-protonated C:CH^+^ base pairs. These structures are most stable in moderately acidic conditions, although certain factors, such as molecular crowding and negative supercoiling, were reported to stabilize them at near-physiological pH (2,3). Thus, the conformational status of any given sequence clearly depends on the specific arrangement of nucleotides, but also on environmental conditions (e.g., pH, solvent, salinity, molecular crowding agents) and responds to external stimuli. Notably, these phenomena are not limited to the *in vitro* space. In fact, the existence of non-canonical secondary structures *in vivo* is supported by a large body of immunochemical, biochemical and biophysical evidence, which has been growing over the last decade. Although a full understanding of these phenomena is still missing, it is generally accepted that such structures act as regulators of genetic and epigenetic transactions (2,4–7). Numerous enzymatic partners interacting with these structures and involved in their homeostasis, either as chaperons or by promoting their unfolding, have been identified (8,9). In fact, imbalances in the delicate equilibria among the different DNA conformations seem to trigger the development of different pathologies such as cancer, infectious, and neurodegenerative diseases (10–13).

Despite the clear relevance of the structural polymorphism of nucleic acids, its high-throughput investigation and, as a consequence, the identification of novel secondary structure-forming sequences is hampered by current methodological limitations. On the one hand, high-resolution methods such as NMR and X-ray crystallography are expensive and time-consuming, despite being highly informative. On the other hand, low-resolution biophysical techniques have reduced costs, but also lack the possibility of a high-throughput implementation. This is perfectly illustrated by the example of circular dichroism (CD), the benchmark technique for this scope. Despite providing a good compromise between information content (14,15), especially when coupled to multivariate (i.e., chemometric) analysis (16,17), and experimental cost, no CD-based high-throughput screening has been reported, to date. In this sense, fluorescence-based methods constitute a valuable alternative. The combination of their high-throughput potential with the decoding power of chemometrics constitutes an important step forward in the quest for a rapid screening method for DNA structures. Recently, we reported a sensor array of fluorescent dyes designed for this scope, providing a proof-of-concept of its applicability for the typing of secondary structures of DNA oligonucleotides (18). However, the use of non-covalent reporter dyes, as in the case of covalently attached labels, can potentially introduce an intrinsic bias. Indeed, covalent or non-covalent interactions of fluorophores with nucleic acids may lead to modification, or induction, of secondary structures of the latter (19,20), engendering a skewing of the screening results. For this reason, the implementation of a label-free method with the same throughput would be an invaluable asset.

In this context, we reasoned that the intrinsic fluorescence emitted by nucleic acids might be exploited for the scope. Similar to isolated nucleotides, single- and double-strands are known to emit in the near-UV spectral range with a low quantum yield (10^−5^ to ~10^−4^); however, certain secondary structures display strongly enhanced fluorescence properties. Thus, the remarkable fluorescent properties of G4 structures were documented about ten years ago (21–23). Upon folding in highly saline buffers, they display a broad emission band, typically peaking between 330 and 420 nm (λ_ex_ = 255–270 nm) (23–25). Most importantly, the quantum yield of G4s (*Φ* = 2.9–3.5 × 10^−4^) is at least 3-fold higher than that of the corresponding single strands (22,23), although these values vary greatly depending on the sequence and other factors. It was proposed that this behavior arises from the interactions of guanine residues in the excited state, including the formation of guanine excimers (26,27). An even stronger emission is observed in some particular cases (e.g., 5′-d[(G_3_T)_3_G_3_]-3′ sequence, hereafter referred to as *G*_*3*_*T*, and its variants, displaying *Φ* values of up to 2 × 10^−3^, i.e., almost 6-fold more fluorescent than other G4s) (25,28), although the reason of such behavior is still a matter of debate. The long-lived red emitting state (λ_em_ = 380–390 nm) has been ascribed to the stacking of guanine bases in defined orientations in the inner part (core) of the G4 structure (25,28,29), or to the formation of excimers of external G-tetrads in 5′–5′ stacked dimeric structures (24,30). Over the years, the effects of guanine base orientation, size and nature of loops, presence and nature of bulges have been investigated, contributing to establish some correlations between the G4 conformation and the emissive properties (29,31,32). Very recently, the investigation on intrinsic fluorescence has been extended to iM structures (33): upon excitation at 267 or 300 nm, iMs display a long-lived, broad emission band centered around 410–420 nm (*Φ* = 3.4–14 × 10^−4^, depending on the specific iM sequence, pH, and λ_ex_). The emission is postulated to arise from the stacking of C:CH^+^ base pairs from the two interpenetrated duplexes (33).

Based on these premises, we hypothesized that the features of steady-state emission spectra (e.g., intensity, maxima, and bandshape) could be exploited to assess the secondary structures adopted by DNA in various conditions. Using a systematic assessment and multivariate analysis of emission spectra of a wide panel of synthetic oligonucleotides with established conformations (G4, iM, single and double strands), we were able to confidently discriminate between different structural motifs. To the best of our knowledge, this is the first report of a large-scale fluorescence-based analysis of DNA sequences to assess their folding preferences, encompassing more than one secondary structure. The results of this work can be exploited for the implementation of a label-free and high-throughput test that could be further extended to genome-wide analyses.

## MATERIALS AND METHODS

### Oligonucleotides and buffer solutions

All chemicals were obtained from Sigma–Aldrich and used as supplied, without further purification. Experiments were performed in aqueous buffers containing 0.01 M lithium cacodylate and 0.1 M of the relevant chloride salt (KCl for buffers A and B, NaCl for buffer C, Table 1), unless stated otherwise. The pH was adjusted to selected values by addition of 0.1 M LiOH solution. Oligonucleotides (sequences: Table S1) were purchased from Eurogentec (RP-Gold Cartridge purification grade) and used without further purification. Stock solutions of oligonucleotides with strand concentration of 100 μM (except for *46AG*: 50 μM and *DDD*: 200 μM) were prepared in deionized water and stored at −20 °C. Samples for CD and fluorescence experiments (referred to as ‘working solutions’) were prepared by diluting the stock solutions with relevant buffers to a concentration of 5.7 μM (except for *46AG*: 2.85 μM and *DDD*: 11.4 μM). Heteroduplexes (Table S1) were prepared by mixing equal volumes of working solutions of the corresponding single strands, giving heteroduplex concentration of 2.85 μM, accounting for the doubled number of nucleotides. Calf thymus DNA (*ct DNA*, Invitrogen, 10 mg mL^−1^) was diluted with deionized water to *c* ≈ 4.2 mM (nucleotides), and further diluted with the relevant buffer to 125 μM so as to obtain a working solution with a comparable nucleotide concentration as in oligonucleotide samples (considering 22 as the average length of oligonucleotides in Table S1). Working solutions were subsequently annealed (5 min at 95 °C), let equilibrate to 20 °C overnight, and stored at 4 °C. For experiments involving mixtures of buffers, DNA samples were annealed separately according to the same protocol and, after equilibration, mixed in the relevant quantities. For experiments carried out at increasing KCl concentration, a 5.7 μM sample of *22AG* in 0.01 M lithium cacodylate buffer (pH 7.2) was gradually supplemented with KCl by addition of aliquots of concentrated (0.01, 0.1, or 1 M) KCl solutions. For experiments carried out at increasing pH, a 5.7 μM sample of *EPBC* in buffer B was titrated with aliquots of 1 M LiOH solution. The resulting solutions were left to equilibrate at 20 °C for 10 min before analysis. The spectra obtained in the last two cases were corrected by multiplication by the dilution factor, to account for the dilution effect.

**Table 1.** Buffer solutions used in this work.

| Buffer | Li<sup>+</sup> | K<sup>+</sup> or Na<sup>+</sup> | pH |
| --- | --- | --- | --- |
| A | ≈ 10 mM | K<sup>+</sup>, 100 mM | 7.2 |
| B | ≈ 10 mM | K<sup>+</sup>, 100 mM | 5.5 |
| C | ≈ 10 mM | Na<sup>+</sup>, 100 mM | 7.2 |

### CD spectroscopy

CD spectra were recorded with a Jasco J-1500 spectropolarimeter. Spectra were recorded using working solutions of DNA in pure buffers (A, B or C) or in mixtures of buffers A and C (1:99, 2:98, 5:95, 10:90, 15:85, 20:80, 40:60, 70:30), unless otherwise stated, in quartz cuvettes with rectangular cross-section (path length 1 × 0.4 cm), with the beam passing through a path length of 0.4 cm. Parameters used for spectra acquisition: wavelength range, 210 to 330 nm; scan speed, 50 nm min^−1^; number of averaged scans, 3; data pitch, 0.5 nm; bandwidth, 2 nm; integration time, 1 s; temperature, 22 °C. Spectra were subsequently corrected for the blank. Finally, spectra were converted to molar dichroic absorption ∆*ε* [M^−1^ cm^−1^] = *θ* / (32980 × *c* × *ℓ*), where *θ* is the CD ellipticity in millidegrees (mdeg), *c* is DNA concentration in M, and *ℓ* is the path length in cm.

### Fluorescence emission spectra

Fluorescence excitation and emission spectra were recorded with a HORIBA Jobin–Yvon FluoroMax-3 spectrofluorimeter, in asymmetric quartz cuvettes (path lengths of 1 × 0.4 cm for emission and excitation beams, respectively). For each sample, two spectra were acquired using λ_ex_ = 260 and 300 nm and emission range of 270–510 nm and 310–590 nm, respectively (slit widths: 5 nm for both excitation and emission beams). For both spectra, data pitch was fixed to 1 nm and integration time to 1 s. All spectra were corrected for the blank, recorded with the appropriate buffer. The obtained spectra were then normalized to the 0–1 interval. When original spectra are presented, these are corrected for the inner filter effect, neglecting the re-absorption of the emitted light (Eq.1):

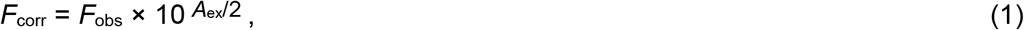

where *F*_corr_ and *F*_obs_ are the corrected and recorded emission, respectively, and *A*_ex_ is sample absorption at the excitation wavelength.

### Single-wavelength absorption measurements

Absorption measurements were performed for all samples at 260, 300 and 400 nm, on a HITACHI U-2900 UV-VIS spectrophotometer. After zeroing of the absorption on a blank sample, the absorption of the sample at the three wavelengths was recorded in a quartz cuvette (1 cm path length). The data at 260 and 300 nm were then used to correct the emission spectra obtained upon excitation at these wavelengths. The absorption at 400 nm was instead used as an internal control.

### Multivariate analysis

Multivariate analysis on the normalized emission spectra (PCA and LDA) was performed with Origin Pro 2018b (OriginLab, Northampton, MA). Data for LDA are presented as Canonical Variables 1 vs. 2 plots, using 85% confidence ellipses. Leave-one-out test was used for internal validation of the LDA method.

### Quantum yield measurements

An appropriate aliquot of the DNA working solution was diluted to 1 mL with buffer A (except for *EPBC* and *i-HRAS2* sequences which were tested in buffer B) so as to obtain a maximum absorbance value of 0.11 at 265 nm. Afterwards, the sample fluorescence was recorded using λ_ex_ = 265 nm and other parameters as described above. The resulting spectrum was integrated between 305 and 505 nm, after blank subtraction. 200 μL of the solution were removed from the cuvette and substituted with 200 μL of the appropriate buffer, and the absorption and emission were measured again. This process was performed four times, to obtain a total of five data points. The integrated areas were then plotted as a function of the absorbance at 265 nm and fitted to a linear model. The same protocol was applied to the reference (quinine sulfate in 0.5 M H_2_SO_4_, *Φ* = 0.546) (34), except that its spectrum was acquired and integrated between 275 and 600 nm. The quantum yield (*Φ*) for each DNA was calculated according to Eq. (2):

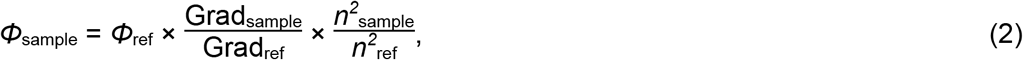

where Grad is the slope calculated from each (Integrated area) vs. absorbance plot, and *n* is the refractive index of the solvent (1.3325 and 1.346 for buffer and 0.5 M H_2_SO_4_, respectively); ‘sample’ and ‘ref’ denote the DNA sample and the quinine sulfate reference, respectively.

## RESULTS

### Intrinsic fluorescence reveals secondary structures of DNA oligonucleotides and their conformational changes

In the first instance, we sought to verify whether steady-state emission spectra could be used to monitor the folding of these structures. First, we focused on the sequence *22AG* (cf. Table S1), a well-studied model G4-forming oligonucleotide (35,36), and proved that its transition from random coil to a G4 structure could be monitored by emission spectroscopy in as much detail as by CD spectroscopy. We recorded CD as well as fluorescence spectra using two excitation wavelengths (λ_ex_ = 260 and 300 nm, as previously used for observing iM fluorescence (33)), in K^+^/Na^+^-free and K^+^-containing buffers (Figure 1). CD spectra of *22AG* show a typical signature (14) of hybrid G4 conformations in the K^+^-containing buffer A, and a spectrum characteristic of a random coil in a K^+^/Na^+^-free buffer (Figure 1, A). The emission spectra mirror this change: in the K^+^-containing buffer, *22AG* displays a strong emission band peaking at 350 nm (similar features were observed with a related telomeric sequence (23)), while in K^+^/Na^+^-free conditions the emission drops to about a half of the intensity and the broad maximum red-shifts to 395 nm (Figure 1, B). The spectra acquired upon 300 nm excitation also display some differences: in the K^+^-containing buffer the emission spectrum peaks at 356 nm, with a large shoulder around 410 nm. In the K^+^-free buffer the maximum is red-shifted to 400 nm, and is slightly less intense (Figure 1, C). Interestingly, comparable results were observed for numerous other G4-forming sequences (Figure S1).

**Figure 1.**
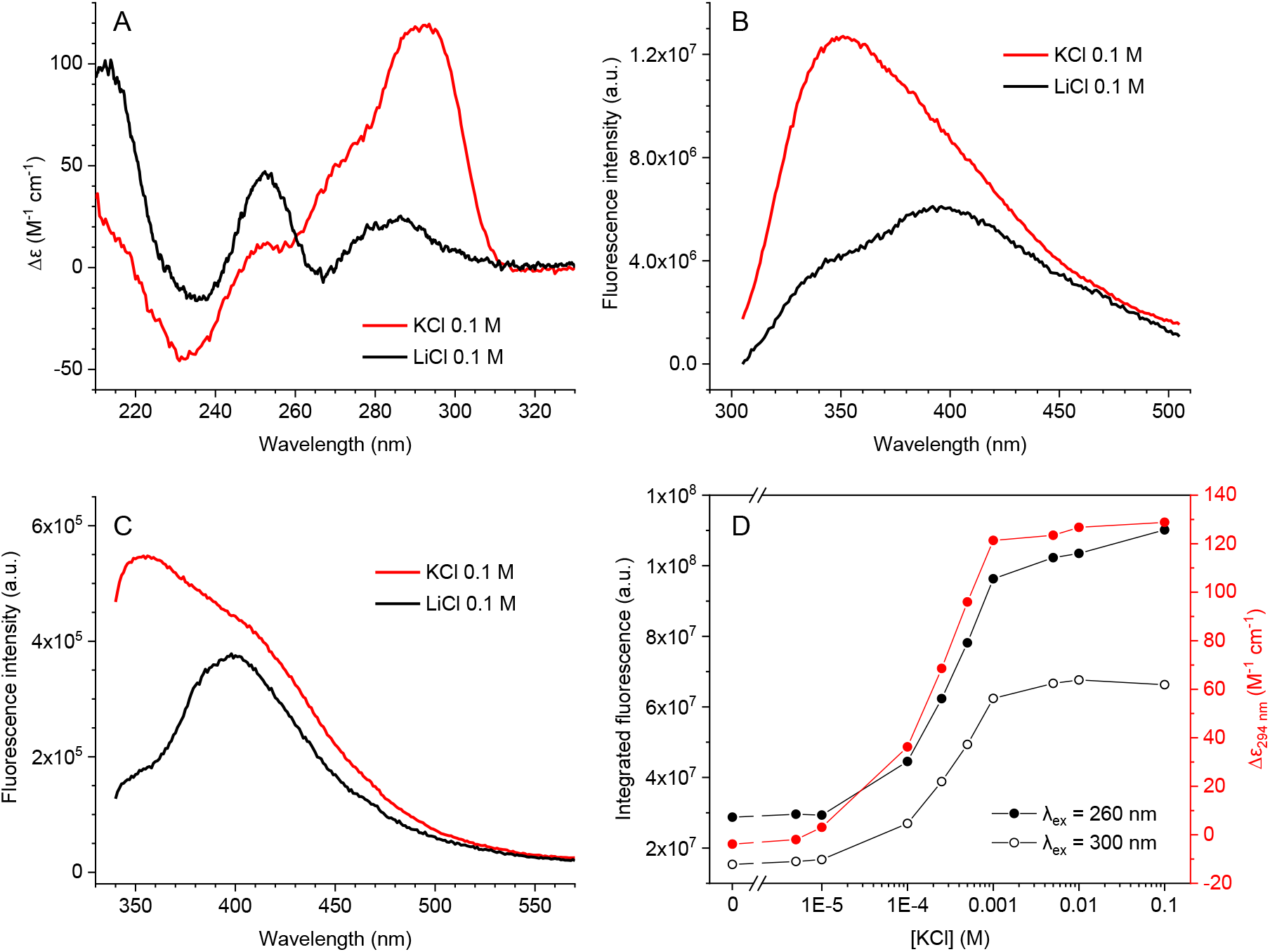
Following the folding of *22AG* by CD and intrinsic fluorescence. A) CD spectra of *22AG* (*c* = 5.7 μM) in K^+^-rich (buffer A) and Li^+^-rich buffers (0.1 M LiCl, 0.01 M lithium cacodylate, pH 7.2). B–C) Fluorescence spectra (B: λ_ex_ = 260 nm, C: λ_ex_ = 300 nm) of the samples presented above. D) Comparison of the integrated fluorescence emission intensity and (red) molar dichroic absorption at 294 nm (from CD spectra) of 5.7 μM *22AG* solutions in 0.01 M lithium cacodylate buffer with variable KCl content (0, 5 × 10^−6^, 1 × 10^−5^, 1 × 10^−4^, 2.5 × 10^−4^, 5 × 10^−4^, 1 × 10^−3^, 5 × 10^−3^, 1 × 10^−2^, and 1 × 10^−1^ M), pH 7.2. Fluorescence spectra were corrected for the inner filter effect.

The descriptive power of intrinsic emission is not limited to the characterization of end-point conditions. Thus, upon progressive increase of K^+^ concentration, *22AG* undergoes a gradual folding (37,38), which is directly reflected in both emission and CD spectra (Figure 1, D; cf. Figure S2 for the full CD and emission spectra). Interestingly, the comparison of characteristic parameters for the two sets of spectra (i.e., the integral fluorescence for emission spectra recorded upon excitation at 260 or 300 nm, and the molar dichroic absorption at 294 nm for CD spectra) leads to overlapping transition profiles, demonstrating the sensitivity of intrinsic fluorescence to the conformation changes accompanying the folding of a G4 structure.

Next, we assessed whether intrinsic fluorescence could be used to monitor more subtle conformation changes. As reported in the literature, *22AG* sequence shifts from a major anti-parallel conformation in Na^+^-rich conditions to a mixture of hybrid forms in K^+^-containing buffers (36,37). This transition can be monitored by recording CD spectra of solutions containing a variable proportion of the two cations: upon increasing the K^+^ / Na^+^ ratio, the CD spectrum gradually switches from the one typical of an anti-parallel G4 (positive maxima at 240 and 295 nm, negative maximum at 265 nm) to that of a hybrid G4 (positive maximum at 290 nm, with two positive shoulders at 265 and 250 nm) (Figure 2, A). Interestingly, an analogous transition is observed in the corresponding fluorescence spectra (Figure 2, B–C). Upon increasing K^+^ / Na^+^ ratio, the emission decreases by about one third of its initial value and the emission maximum blue-shifts from 375 to 350 nm. The information provided by the two methods is perfectly aligned, as shown in Figure 3, D (cf. Figure S3 for the analysis of fluorescence spectra obtained with λ_ex_ = 300 nm).

**Figure 2.**
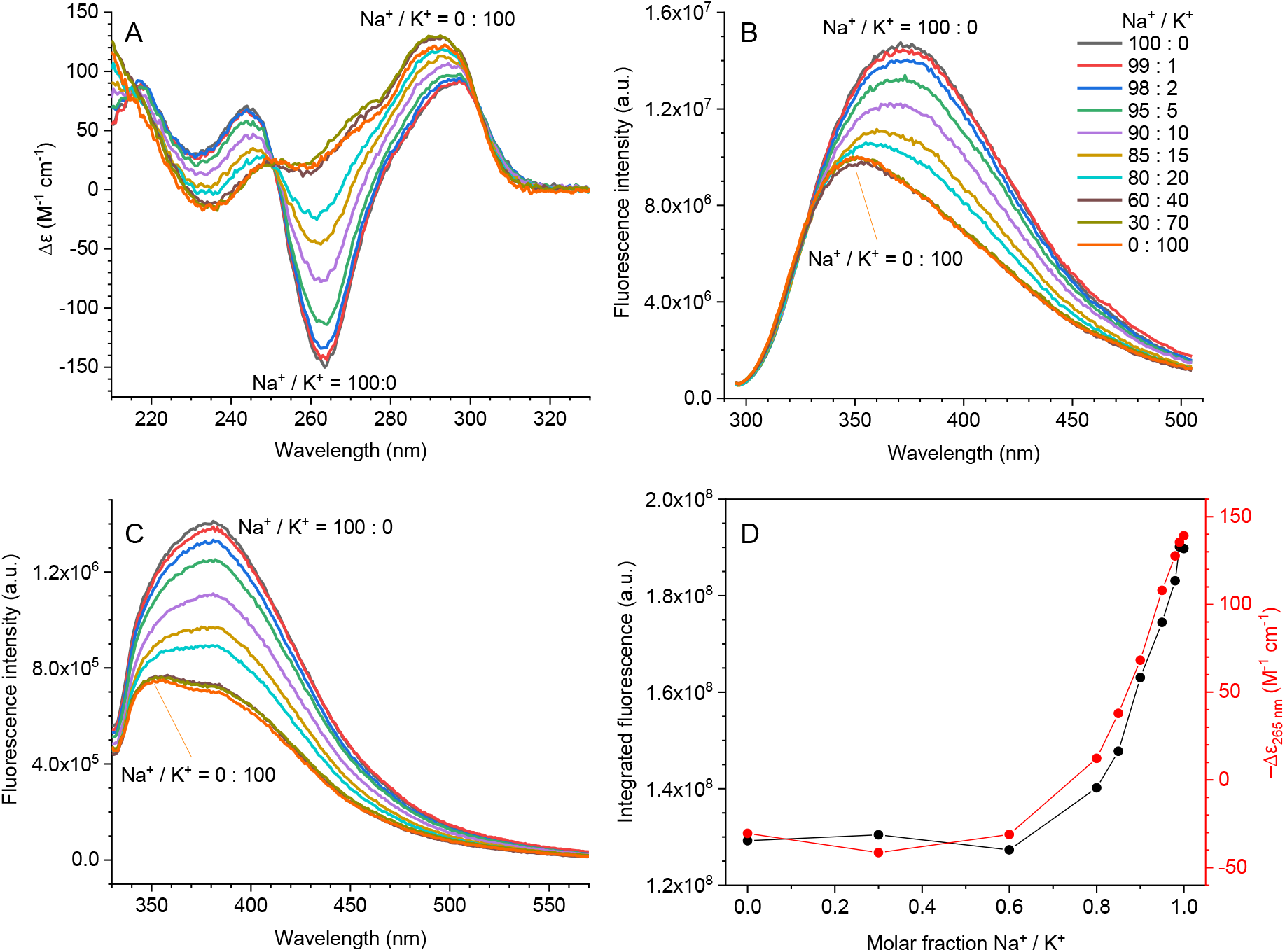
Conformational transition of *22AG* from anti-parallel to hybrid G4 monitored by CD and intrinsic fluorescence. A) CD spectra of *22AG* solutions (*c* = 5.7 μM) containing variable proportions of Na^+^ and K^+^ (total NaCl + KCl concentration of 0.1 M in all solutions, lithium cacodylate buffer 0.01 M, pH 7.2). B–C) Corresponding emission spectra (B: λ_ex_ = 260 nm, C: λ_ex_ = 300 nm). D) Comparison of the integrated fluorescence intensity (λ_ex_ = 260 nm) and molar dichroic absorption at 265 nm (from CD spectra) at the various KCl / NaCl ratios.

**Figure 3.**
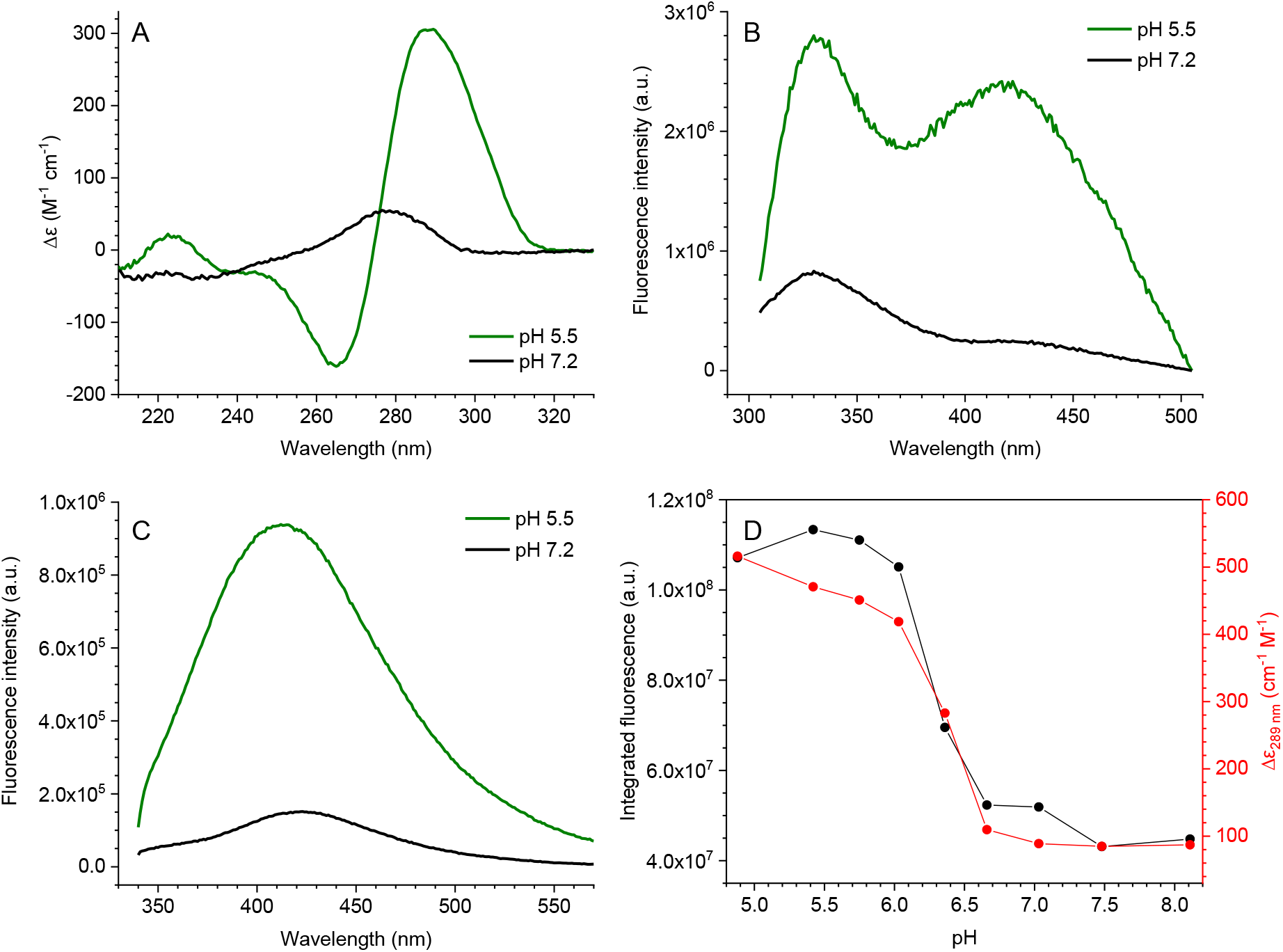
Following the folding of *EPBC* to iM structure by CD and intrinsic fluorescence. A) CD spectra of *EPBC* (*c* = 5.7 μM) in neutral and acidic conditions (buffers A and B, respectively). B–C) Fluorescence spectra (B: λ_ex_ = 260 nm, C: λ_ex_ = 300 nm) of the samples presented above. D) Comparison of the integrated fluorescence emission (λ_ex_ = 260 nm) and (red) molar dichroic absorption at 289 nm (from CD spectra) of 5.7 μM *EPBC* solutions at variable pH (4.9–8.1, 0.01 M lithium cacodylate buffer, 0.1 M KCl). Fluorescence spectra were corrected for the inner filter effect.

Finally, we studied the transition of *EPBC* from random-coil to iM structure. In this case, the switch from neutral to acidic pH is accompanied by the formation of a strong positive CD band at 289 nm and a moderately intense negative band at 265 nm (Figure 3, A). In the emission spectra obtained upon 260-nm excitation, the intensity of the 330-nm maximum increases about 3-fold and a second broad emission band appears, centered at around 420 nm (Figure 3, B). The emission spectra obtained upon excitation at 300 nm vary accordingly: the emission maximum (around 410 nm) at pH 5.5 is about five times more intense than at neutral pH (Figure 3, C). A similar behavior was observed for other iM-forming oligonucleotides (Figure S4). As previously observed for the folding of *22AG*, the gradual conformational transition can also be monitored for *EPBC*. In fact, the unfolding of the iM structure upon increase of pH is directly reflected in both CD and emission spectra (Figure 3, D; cf. Figure S5, A–B for the full spectra), and the comparison of the characteristic parameters for the two sets of spectra (i.e., integrated emission intensity and the molar dichroic absorption at a characteristic wavelength, from CD spectra) leads to overlapping transition profiles. Similar trends were observed when using the data from emission spectra obtained upon excitation at 300 nm (Figure S5, C–D).

### Emission properties of various DNA sequences

Based on these promising data, we assessed the potential of intrinsic fluorescence as a reporter of oligonucleotide conformation on a larger scale. Towards this end, we screened an 89-membered set of DNA analytes with different and well-established conformations (Table S1). Specifically, the selected sequences comprise 46 G4 structures (including 14 hybrid, 18 parallel, and 14 anti-parallel folds), 14 iM-forming sequences (iMFS) of varying stability, 14 duplexes (including three auto-complementary sequences, three hairpins, three hetero-duplexes, and one highly polymerized genomic DNA), and 12 single-strands with different base composition and purine vs. pyrimidine content. For the sake of homogeneity of our assay, we deliberately omitted bi- and tetramolecular G4 structures (such as d[TG_4_T]_4_ and d[G_3_T_4_G_3_]_2_), as well as unimolecular G4 structures with a known propensity to form dimers or higher aggregates, such as *N-myc* (39), *93del* (40), *G*_*3*_*T* and its analogues (41). The conformations adopted by all sequences were initially assessed by CD spectroscopy in three different buffers (A, B and C). The results (Figures S6 and S7, summarized in Table S2) confirmed that most putative G4-forming sequences adopted the expected conformation in buffer A (Figure S6, A/D/G), with the exception of *LWDLN-1*, *19wt* and *SP-PGQ3* whose CD spectra did not agree with the reported conformations, most likely due to the differences in experimental conditions employed for their structural characterization (cf. Table S2 footnote). Subtle or no changes were observed when these sequences were reassessed in K^+^-containing acidic conditions (buffer B). All putative G4-forming sequences appear to be folded under these conditions, and most maintained the same conformation as in buffer A, except for *Bcl2Mid* which underwent a conformational change from hybrid to parallel form, and *hras-1* whose CD spectrum gave evidence of a partial conformational change (Figure S6, B/E/H, and Table S2). The use of the Na^+^-containing buffer C was, instead, more problematic, since 12 out of 46 sequences were completely or partially unfolded under these conditions). In addition, 16 out of 46 sequences underwent considerable conformational changes, most typically a change from hybrid (*22AG* as described above, *46AG*, *26TTA*, *23TAG*, *24TTA*, *chl1*, *UpsB-Q-3*) or parallel (*c-kit2-T12T21*, *VEGF*, *Myc1245*) to anti-parallel forms (Figure S6, C/F/I, and Table S2). Therefore, we decided to discard buffer C for the following emission studies. With regard to iMFS, these appeared mostly folded in buffer B, as expected in slightly acidic conditions. In buffer A, all iMFS display CD spectra compatible with a random-coil arrangement, likely due to insufficient protonation, although the presence of a small fraction of folded structure cannot be ruled out on the basis of CD spectra (Figure S7, A–B). Finally, according to their CD spectra, double-stranded and single-stranded sequences adopted the expected conformation in all buffers, regardless of the specific pH (Figure S7, C–F).

Next, we recorded emission spectra of all aforementioned sequences in buffers A and B, using excitation at 260 and 300 nm (Figures S8 and S9, respectively). The features of the emission spectra (i.e., band shape and intensity) were found to vary to large extents inside each group. However, several patterns could be identified upon naked-eye examination of the spectra. On the overall, the emission intensities were higher upon excitation at 260 nm than at 300 nm, due to the differences in sample absorbances at these wavelengths. As a general behavior, the intrinsic emission of G4 structures is significantly more intense (roughly 3–4-fold, based on the comparison of the integrated fluorescence intensities) than that of other structures upon 260-nm excitation (Figure 4, A–B). This difference is more limited upon excitation at 300 nm (G4 emission is only 1.5-fold higher than that of other structures, Figure 4, C–D). Interestingly, iM structures become the most fluorescent in buffer B upon 300-nm excitation, with a median intensity 1.5-fold higher than that of G4 structures under the same conditions (Figure 4, B). This difference is not observed in the data obtained upon excitation at 260 nm, due to the considerable spectral shape changes. In fact, in this case, the appearance of the iM-characteristic, broad peak centered around 425 nm in buffer B is accompanied by a reduction of the 320-nm maximum by approximately a half (Figure S8, G–H), leading to the decrease of the integrated emission intensity (Figure 4, B). On the contrary, the iM-characteristic red-shifted peak is predominant in the spectra obtained upon excitation at 300 nm, with an intensity increase of about 3.5-fold upon moving from buffer A to B (Figure S9, G–H). This is likely due to the fact that a longer wavelength selectively excites the iM-characteristic band that otherwise appears only as a shoulder.

**Figure 4.**
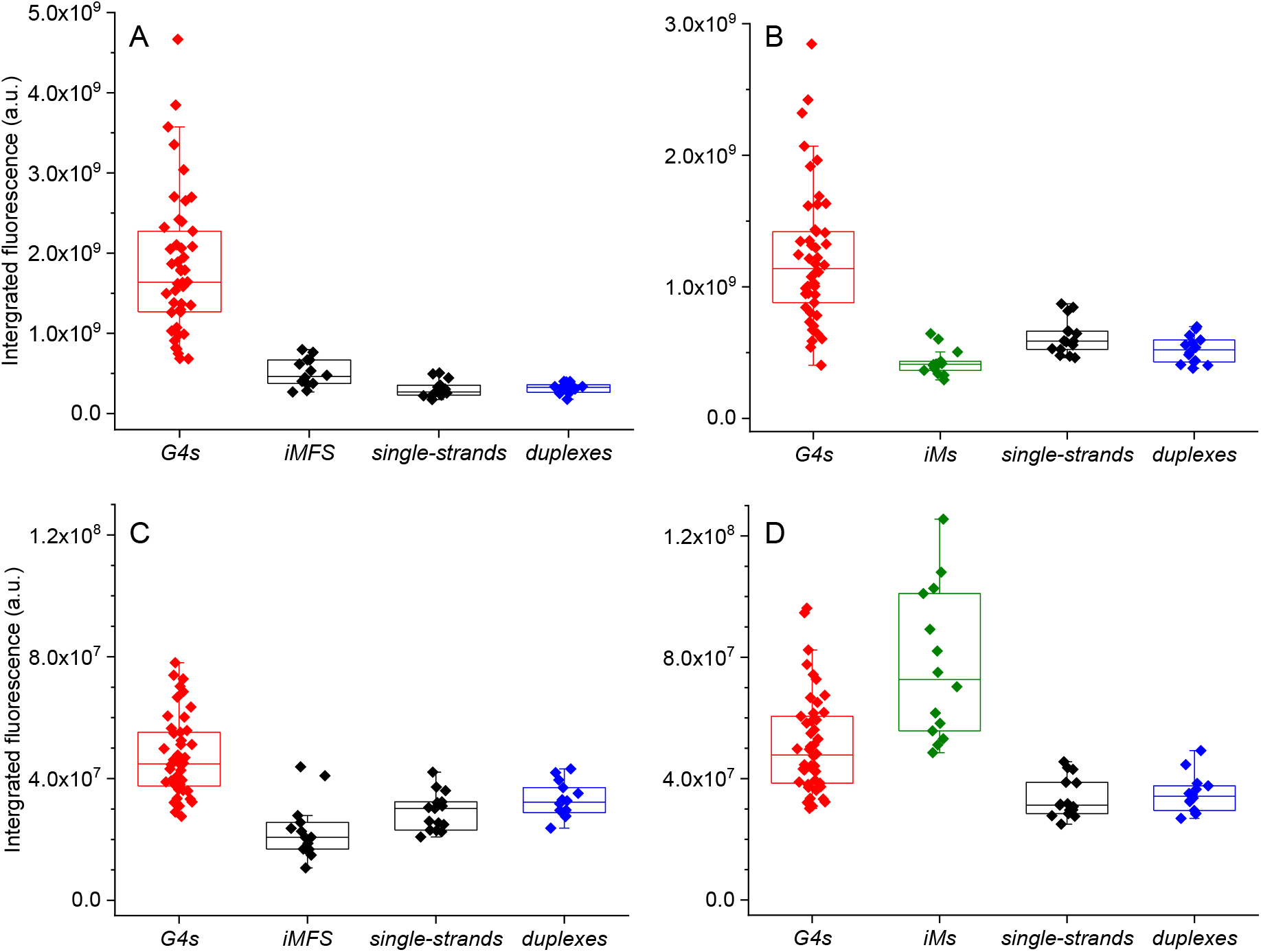
Box plots of the integrated fluorescence intensity of DNA samples grouped according to their conformation. A–B) λ_ex_ = 260 nm in A) buffer A and B) buffer B; C–D) λ_ex_ = 300 nm in C) buffer A and D) buffer B.

The spectral shape was also found to be informative of the oligonucleotide conformation, especially considering the spectra obtained upon 260-nm excitation, as demonstrated by the comparison of group-averaged normalized emission spectra (Figure 5). In addition, the use of normalized emission spectra avoids the differences in intensity arising from the differences in the length of oligonucleotides (and thus in their absorbance at λ_ex_). As shown in Figure 5, A–C, and Figure S10, A–C, G4 structures give broader emission spectra in both buffers A and B, with maxima centered between 330 and 350 nm. The spectra are quite broad, loosing gradually in intensity at longer wavelengths. No general trends are observed that could enable the discrimination between different G4 topologies. In contrast, single- and double-stranded structures display significantly sharper spectra in both conditions, peaking around 350 and 330 nm, respectively (Figure 5, D–E and Figure S10, D–E). In all cases, a more or less pronounced shoulder is observed between 400 and 500 nm. The same is true for iM-forming sequences in buffer A (Figure 5, F), in which they are mostly unfolded. However, at lower pH (buffer B), a characteristic shoulder appears between 375 and 475 nm, whereas the maximum remains fixed at around 330 nm (Figure 5, G). The spectra obtained using λ_ex_ = 300 nm do not display the same degree of shape variability (Figure S11). In this case, most sequences display a maximum between 310 and 330 nm and no shoulders in any of the buffers, regardless of the adopted conformation. The only remarkable difference is observed for some G4 structures, displaying a blue-shifted shoulder of varying intensity.

**Figure 5.**
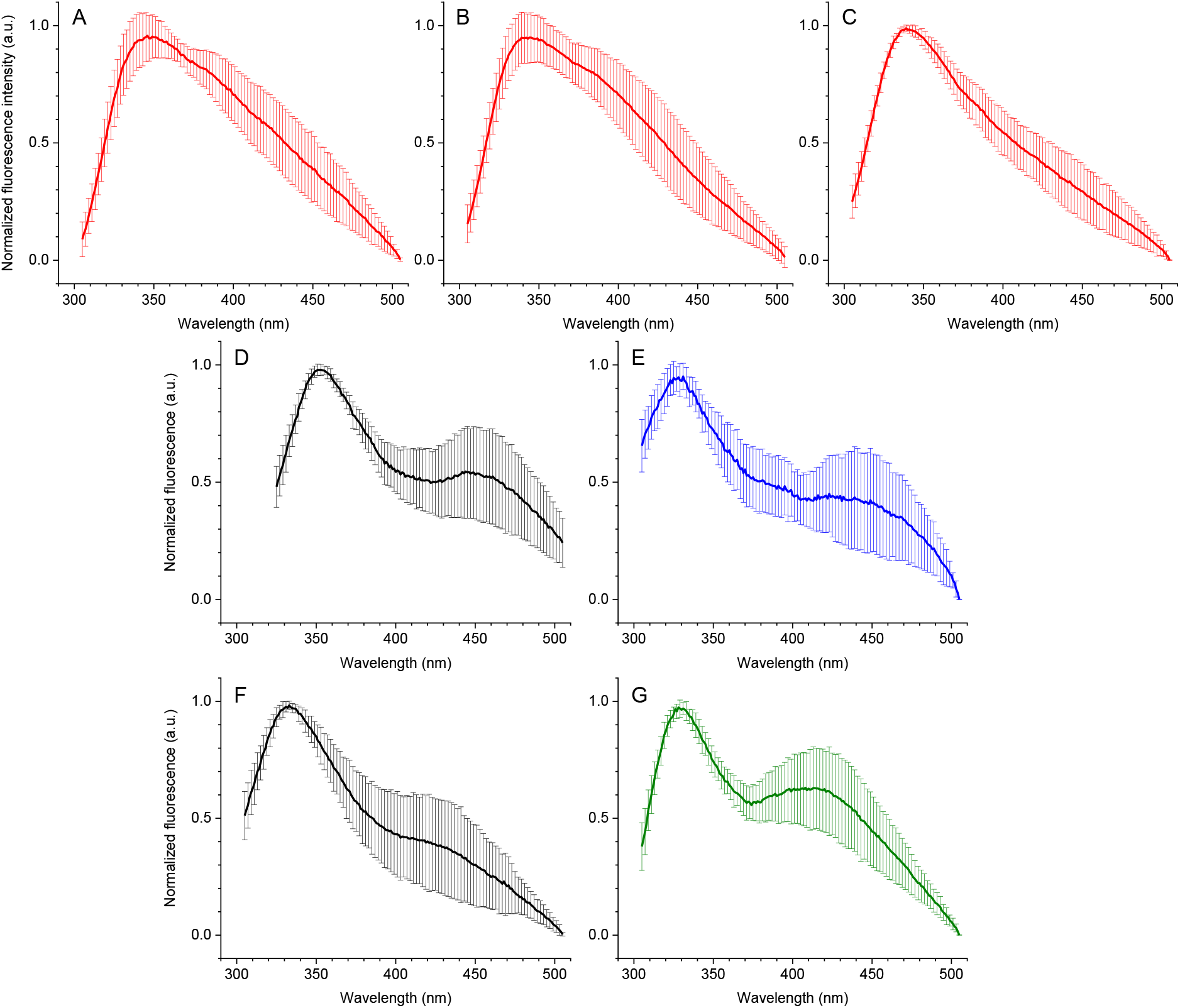
Group-averaged, normalized emission spectra (λ_ex_ = 260 nm, buffer A, unless stated otherwise) of the tested oligonucleotides, grouped according to their conformation (as confirmed by CD spectra). A) hybrid G4s; B) parallel G4s, C) anti-parallel G4s, D) single strands, E) duplexes, F) iM-forming sequences, G) the same as F in buffer B. Error bars represent the standard deviation of the emission at a given wavelength inside each conformational group. *LWDLN-1*, *19wt* and *SP-PGQ3* were excluded from the analysis.

### Principal component analysis of the emission spectral dataset

To extract more information, the emission spectral dataset was subjected to principal component analysis (PCA). PCA is an unsupervised multivariate method, which describes the variance of the examined data matrix through a reduced number of variables and highlights the similarities between the analytes displaying similar response patterns. Thus, four emission spectra generated for each analyte (corresponding to two excitation wavelengths and two buffers, A and B) would account for 894 variables if the information obtained at each wavelength (1 nm data pitch) were to be used. These would be reduced to 447 variables for the spectra obtained in just one of the buffers, or to 201 (or 246) variables for a single spectrum (corresponding to the emission spectral windows). PCA analysis enables to combine these data into a limited number of principal components (PCs) and facilitates their interpretation.

In order to understand which spectra were more suitable for the analysis, we first performed a preliminary PCA on each separate set of spectra acquired with different excitation and buffer conditions. Remarkably, upon using normalized spectra obtained in buffer A upon 260 nm excitation, we readily observed a good separation between G4 structures, mostly occupying the upper part of the PCA plot, and all other conformations (Figure 6, A). In this case, the putative iM-forming sequences (iMFS) were combined with the group of single-stranded oligonucleotides, since they are mostly unfolded under these conditions. No clear difference could be observed between single- and double strands, in agreement with what could be qualitatively inferred from inspection of the spectra. When the same analysis was run on the data obtained in buffer B under the same excitation wavelength conditions, a partial separation could be inferred for the i-motif structures: these sequences form a broad cluster overlapping with those of G4s and single- and double-strands (Figure 6, B). In both buffers, the spectra obtained upon excitation at 300 nm were less suitable for the analysis (Figure S12). In fact, in both cases the separation between the groups was less efficient due to the little amount of variance described by PC2 (18.4% and 16.4% for buffers A and B, respectively).

**Figure 6.**
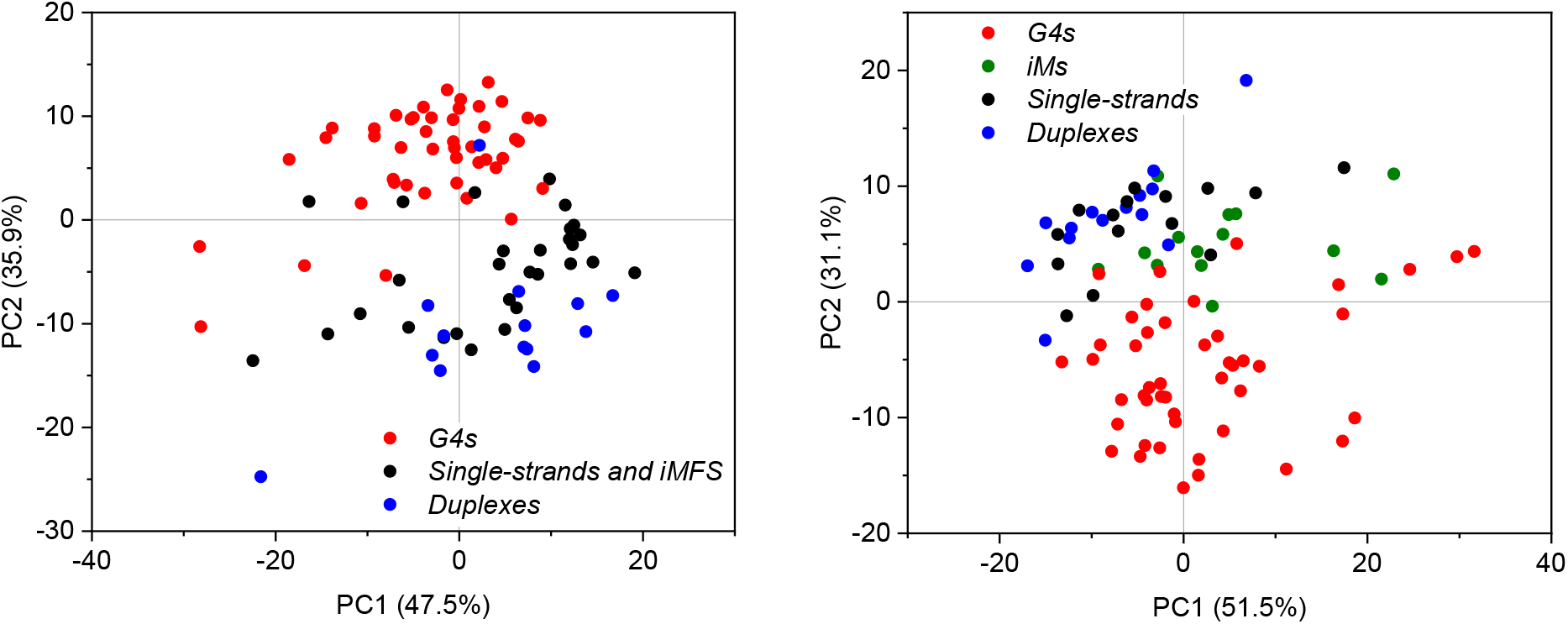
PC1 vs. PC2 plots resulting from the PCA of normalized emission spectra recorded A) in buffer A (λ_ex_ = 260 nm); B) in buffer B (λ_ex_ = 260 nm). In (A), the putative iM-forming sequences (iMFS) sequences are grouped together with single-strands (in black), as they are mostly not folded in this buffer. In (B), they are shown in green, since they adopt a distinct conformation.

In order to improve the clustering of iM structures and thus be able to detect the three separate groups (i.e., G4s, iMs and ‘others’, from now on used to designate the group of single-and double-strands) in a single plot, we attempted PCA on a combined dataset, including the spectra obtained upon 260-nm excitation in both buffers A and B. We speculated that, since the data obtained in each buffer alone were not sufficient to reveal the cluster of iM structures, a change in fluorescence properties between the buffers A and B, specific for iM-forming sequences, could provide an additional information element. Indeed, the visualization and separation of the three groups was significantly better in this case, especially when using the first three principal components (Figure 7, A). The three principal components presented in the plot account for 82.3% of the dataset variance. Inspection of the scree plot (Figure S13) shows that five components out of twenty that had been calculated are actually significant and describe up to 92.3% of the dataset variance. In order to better understand the concrete meaning of each of the PCs, we examined the various 2D plots (PC1 vs. PC2, PC1 vs. PC3 and PC2 vs. PC3: Figure 7, B–D). For PC2 and PC3, we could infer a clear correlation with the propensity of samples to adopt a G4 or iM structure, respectively. In fact, G4 structures have generally positive PC2 scores (‘*G4-likeness*’), whereas iMs, single- and double-stranded sequences have negative ones (Figure 7, B and D). At the same time, iMs are the only structures with high negative value scores for PC3 (‘*iM-likeness*’), whereas all the other analytes display values around zero (Figure 7, C and D). With respect to PC1, data points are distributed quite evenly along the axis, suggesting that the correlation is not purely conformational.

**Figure 7.**
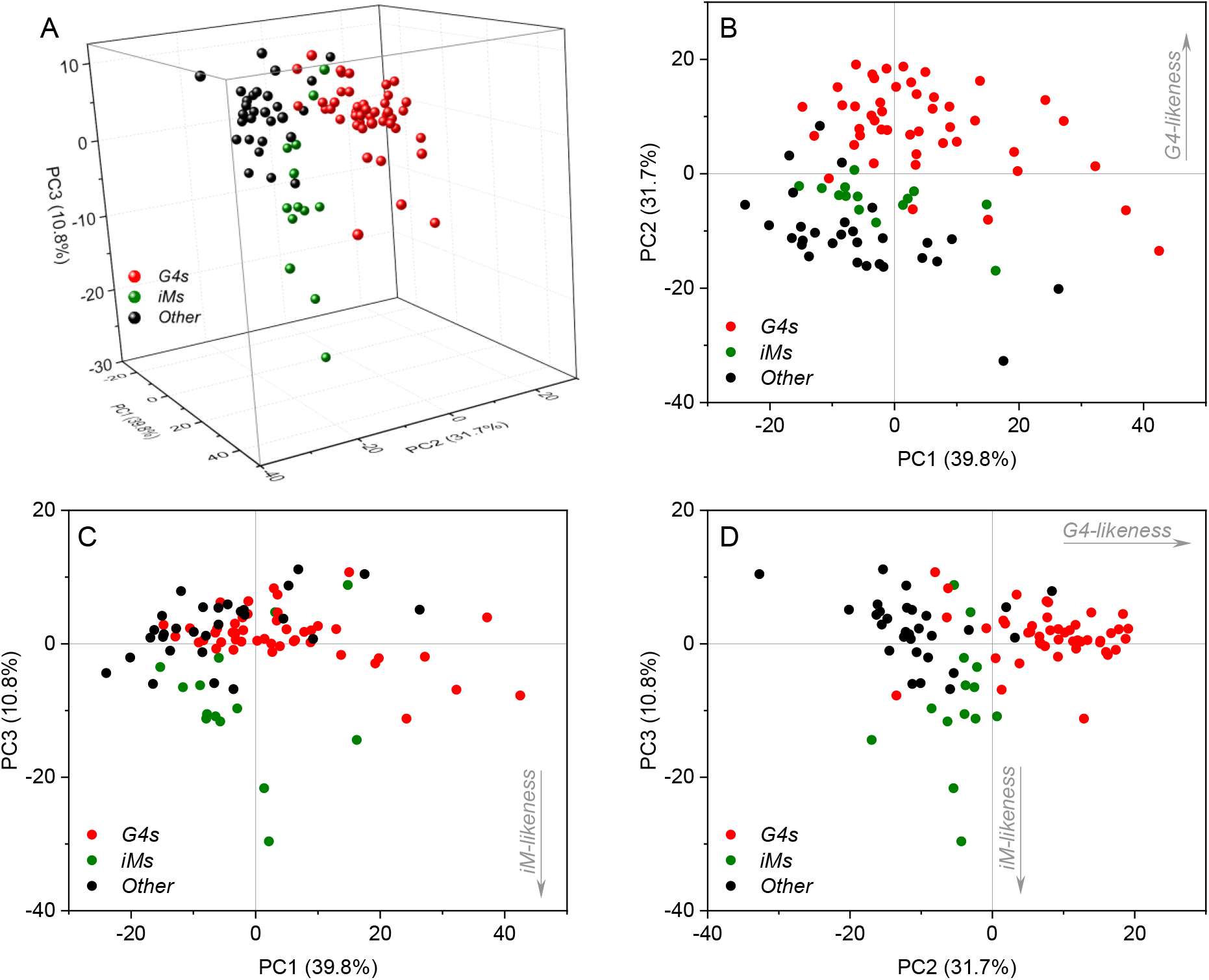
PCA of the combination of normalized emission spectra recorded in buffers A and B (λ_ex_ = 260 nm). A) 3D PCA plot (PC1 vs. PC2 vs. PC3); B–D) corresponding 2D plots.

### Dataset reduction and linear discriminant analysis of emission spectra

As a next step, we attempted to reduce the number of variables (and thus the dataset size) by selecting spectral regions that would be most relevant for sample grouping. Ideally, these should correspond to the wavelengths at which the analytes belonging to the same conformational group share similar spectral shapes, whereas the analytes from different groups show significant differences. To identify such zones, we calculated the intra-group variance for the normalized values obtained at each wavelength and, from the resulting three sets of values, the inter-group variance at each wavelength. These were plotted as a function of the wavelength (Figure S14). From this graph, four zones (324–333 nm and 374– 393 nm for the spectra obtained in buffer B; 333–343 nm and 378–388 nm for buffer A) in which the inter-group variance was relatively high were selected for further examination. In this choice, we tried to favor the spectral zones in which the intra-group variance of iM structures was low, since this group showed the poorest clustering. The selected wavelengths account for 52 variables out of the 402 arising from the two combined spectra, corresponding to an 87.5% reduction of the dataset size. PCA was ran on this reduced dataset, to verify the quality of the clustering under these conditions (Figure S15, A), and gave satisfactory results, with the three groups still appearing as relatively well separated.

The resulting dataset was further reduced in order to subject to it to linear discriminant analysis (LDA). This is a supervised multivariate method, enabling to assign a test analyte to one of the *a priori* defined groups on the basis of the similarity of, in our case, emission spectrum profile, with those of the elements of the training set. The limiting factor in LDA is constituted by the group sizes, since the number of variables cannot exceed the number of training samples in each group. In our case, the group of iMs only contains 14 oligonucleotides, thus restricting the number of exploitable variables to 14. We thus evenly selected 14 wavelengths out of the 52 identified ones and run again PCA to ascertain that the information content and the groups separation were retained. Satisfyingly, this was the case (Figure S14, B), and we proceeded to perform LDA on the established dataset (Figure 8).

**Figure 8.**
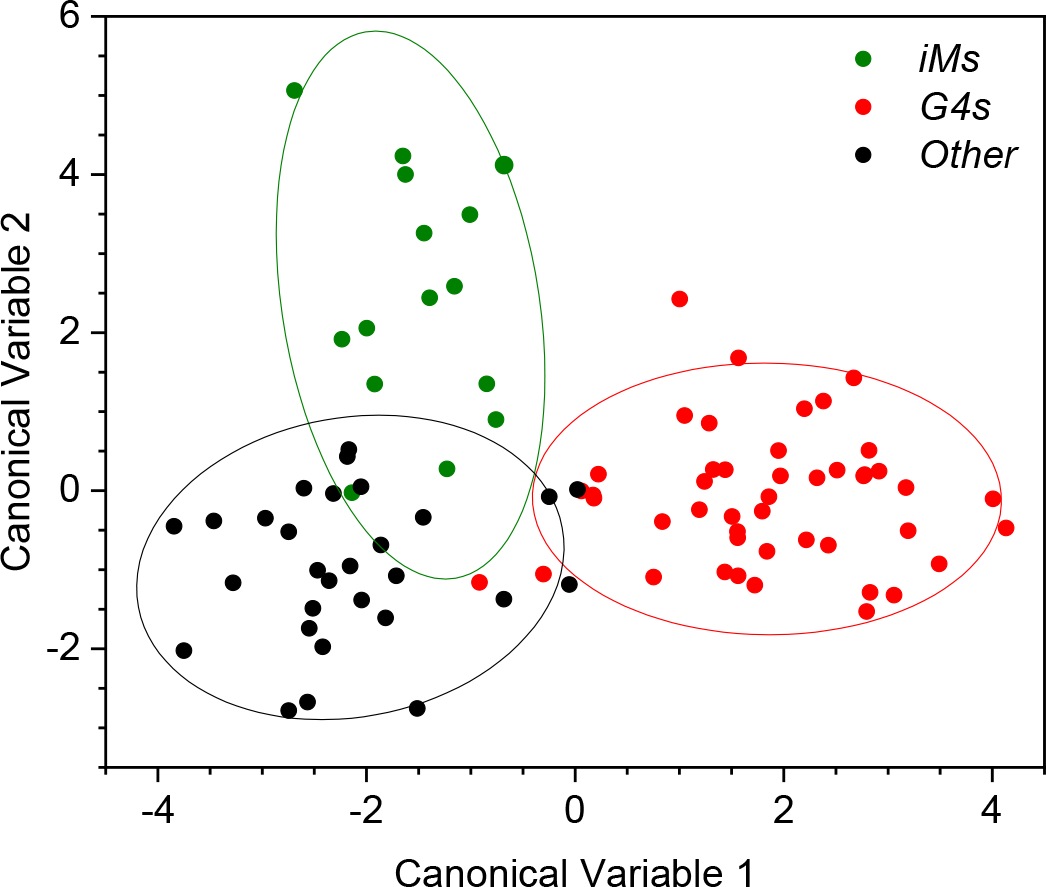
Canonical variable 1 vs. Canonical variable 2 plot obtained from LDA of the reduced emission dataset (normalized emission data recorded at 335, 338, 341, 380, 383 and 386 nm in buffer A, and 329, 332, 376, 379, 382, 385, 388 and 391 nm in buffer B, λ_ex_ = 260 nm) on 89 characterized sequences. Ellipses represent 85% confidence zones for each group.

Remarkably, DNA sequences were found to cluster relatively well, according to their conformation. Although the partial overlap of the clusters could not be avoided (in particular, those of iMs and ‘other’ sequences), the validation by the leave-one-out method confirmed a 90.8% of correct identifications. Considering the complexity of our task with respect to usual chemometrics applications, this error rate is considered quite satisfactory.

### Chemometric assessment and fluorescence properties of novel sequences

In order to assess the behavior of the established multivariate methods upon analysis of sequences with unknown conformation, we recorded the emission spectra of a number of test sequences and analyzed them by PCA and LDA. In the first instance, we analyzed four randomly generated, 24- to 25-mer sequences with moderately high *G4Hunter* scores (*RND-HS1* to *RND-HS4*, Table S3). PCA analysis of their emission spectra revealed that in the PCA plot, *RND-HS1* fell definitely closer to the group of single- and double-strands (‘other’), *RND-HS2* was located in between the two groups, and *RND-HS3* and *RND-HS4* fell closer to the center of the G4 group (Figure S16, A). Subsequent LDA testing confirmed this interpretation (Figure S16, B), assigning *RND-HS3* and *RND-HS4* to the G4 group (with probabilities *P* = 0.98 and 0.80, respectively) and *RND-HS1* and *RND-HS2* to the group of single- and double-strands (Table S3); interestingly, the probability of *RND-HS2* to belong to the G4 group was not negligible (*P* = 0.095). To test the validity of the LDA prediction, we recorded ^1^H NMR, CD and TDS spectra of the four test oligonucleotides. NMR data (Figure S17) showed that all four sequences showed the presence of imino proton peaks characteristic of the formation of secondary structures (42). In the case of *RND-HS1* sharp peaks were observed between 12.5 and 14 ppm, indicative of formation of Watson–Crick base pairs and a duplex-type structure, despite the presence of a small hump at 10–11.5 ppm. *RND-HS2* displayed sharp peaks at 12.8 to 13.2 ppm indicative of a Watson–Crick base pair related structure and a broad peak centered at 10.7 ppm characteristic of Hoogsten base pairings; it is noted that the formation of intermolecular structures could be favored by the high concentration of oligonucleotides required for NMR experiments (130 μM, i.e. 23-fold higher with respect to fluorescence and CD studies). In the case of *RND-HS3*, NMR spectrum in the imino region showed several sharp peaks in 10–12 ppm region and a sharp peak at 13 ppm, characteristic of G4-duplex hybrid structures; indeed, the formation of a hairpin-type structure is possible, considering the presence of two 6-nt complementary runs of GC base pairs in this sequence (Table S3). Finally, *RND-HS4* exclusively displayed broad signals in the Hoogsten base-pair region giving evidence of formation of multiple G4 structures, in agreement with the LDA prediction based on the fluorescence data. The inspection of CD spectra of these sequences in buffer A suggested that *RND-HS1* is unlikely to fold into a G4 structure, whereas the other three sequences might adopt such a conformation (Figure S18, A). Furthermore, the results of thermal difference spectra (43) were in agreement with this assignment: *RND-HS3* and, particularly, *RND-HS4* showed negative bands in the 295−300 nm region, giving evidence of at least partial formation of G4 structures, whereas *RND-HS1* and *RND-HS2* were devoid of this band (Figure S16, B), supporting our NMR data and fluorescence-based LDA typing.

Subsequently, we tested a number of sequences adopting secondary structures other than those represented in the training set. These include the stacked dimeric G4 structure known to be the most fluorescent unmodified oligonucleotide described to date *G*_*3*_*T*) (25,28,41), three left-handed G4 structures (*ZG4*, *Block2∆*, *2xBlock2*) (44,45), three peculiar non-G4 quadruple helices (*VK1*, *VK2*, *VK34*) (46,47), two A-type duplexes (*(G*_3_*C*_3_)_3_ and *(G*_3_*C*_3_)_2_) (48,49), a G-hairpin (*SC11*) (50), and three supposed single strands displaying peculiar CD signatures, namely *ss8*, *24non096* and *scr26* (Table S4). The CD spectra of these sequences in buffers A and B are presented in Figure S18. In the PCA plot based on the fluorescence data (Figure 9, A), most non-G4-forming sequences were distributed in the zone of single- and double-strands, except for *VK1* and *VK2*, which fell in the G4 zone. In agreement with this result, all sequences were assigned to the corresponding groups upon LDA analysis (Figure 9, B, and Table S4), except for *ss8*, which was mis-assigned to the iM group (*P* = 0.87) and *scr26* which was assigned to iM and “other” groups with almost equal probabilities (*P* = 0.48 and 0.51, respectively). This result indicates that the method yields a relatively low rate of false positive results with respect to iM and G4 groups. At the same time, the inconsistency between the group assignment and the expected structure (as can be inferred from the sequence analysis) points out to some peculiarities, for example in the case of *ss8* and *scr26*, as already evidenced by their CD spectra and, in the case of *ss8*, the anomalous results of the fluorescent-probe analysis (51). Such anomalies clearly call for further in-depth studies.

**Figure 9.**
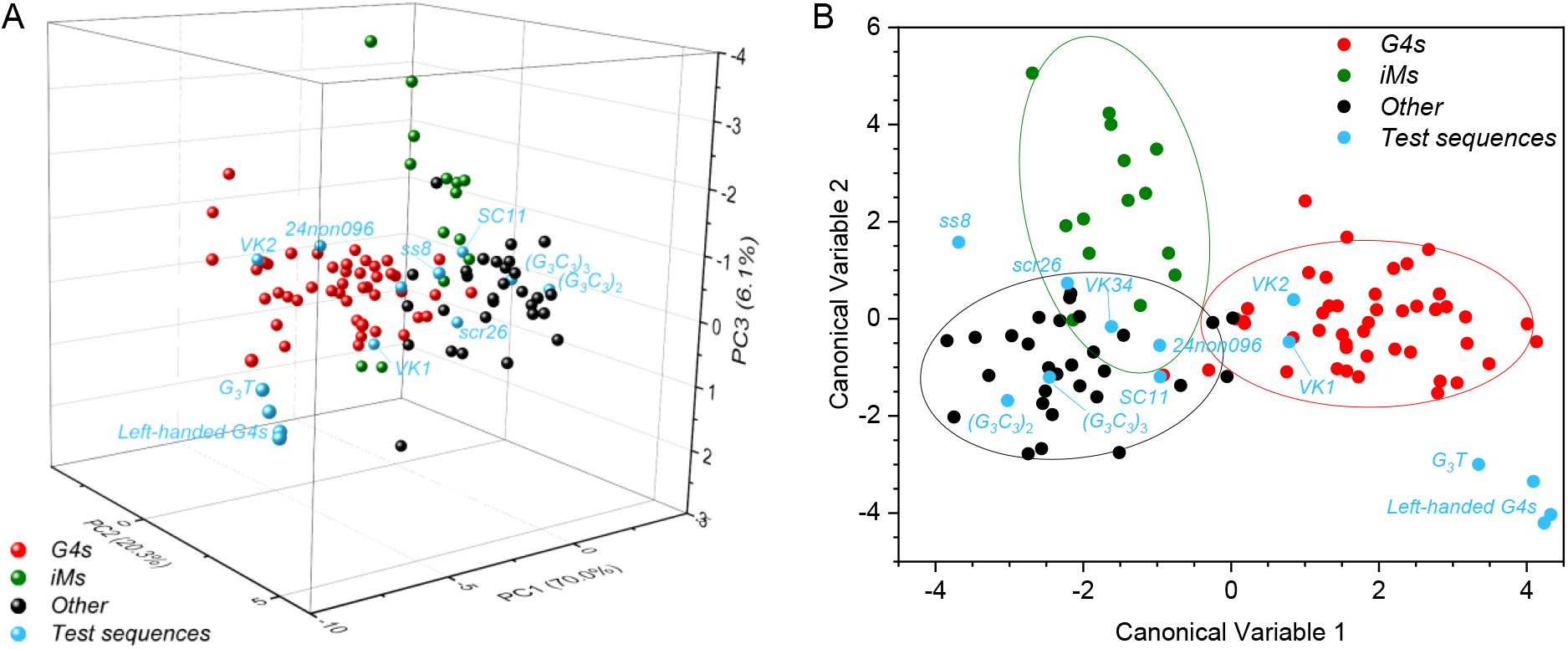
A) PC1 vs. PC2 vs. PC3 plot obtained from the analysis of the training set (reduced dataset), supplemented with the emission data obtained for test sequences with peculiar conformations *G*_*3*_*T*, *ZG4*, *Block2∆*, *2xBlock2, VK1*, *VK2*, *VK34*, *(G*_*3*_*C*_*3*_)_*3*_, *(G*_*3*_*C*_*3*_)_*2*_), *SC11*, *ss8*, *24non096* and *scr26*). B) LDA plot for the same dataset.

Another peculiar observation following from the results of the multivariate analysis is represented by the left-handed G4 structures which, along with *G*_*3*_*T*, occupy an otherwise empty region of the PCA plot (Figure 9, A). Accordingly, they spot in the bottom right corner of the LDA plot, relatively far away from all other clusters (Figure 9, B); nonetheless, LDA unambiguously assigned these structures to the G4 group (Table S4). This behavior is likely due to the peculiar emission spectra of these structures, which are characterized by a sharp emission band peaking around 385 nm in both buffers A and B (Figure S20). This value is 35 to 55 nm red-shifted with respect to the values observed for other G4 structures and, more in general, for all the examined oligonucleotides. Of note, the emission completely fades off upon switching from a K^+^ to a Na^+^ rich buffer, as a result of the structure unfolding evidenced by the corresponding CD spectra (Figure 10). The peculiarities of left-handed G4s are not limited to the shape and position of their fluorescence bands, but also extend to the emission intensities. In fact, upon comparison of the raw emission data in the maxima, left-handed G4s appear to be on average 2.2-fold more fluorescent than other representative G4 sequences. A quantitative description of this phenomenon is offered by the values of fluorescence quantum yield (*Φ*, Table 2; cf. Figure S21 for the determination of these values). Indeed, left-handed G4s prove in general more fluorescent (*Φ* = 2.0 × 10^−3^ to 3.1 × 10^−3^) than other G4 structures (*Φ* = 4.0 × 10^−4^ to 2.4 × 10^−3^) and other sequences (*Φ* = 2.8 × 10^−4^ to 1.4 × 10^−3^). Interestingly, the quantum yields of left-handed G4 structures are almost as high as that of the *G*_*3*_*T* sequence (*Φ* = 3.7 × 10^−3^ in our conditions, i.e., almost two-fold higher than 2 × 10^−3^ as reported in (28)). The reason for this similarly enhanced fluorescence might reside in shared structural features. Indeed, all reported left-handed G4s share a particular stacking mode of G-tetrads, different from the ones observed in most other G4 structures irrespective of their topology (Figure S22). Notably, the same 5′–5′ stacking mode (termed ‘5/6-ring stacking’) is observed in a close analogue of *G*_*3*_*T* (*J19*, PDB: 2LE6) forming a stacked dimer in solution, and was proposed to be at the origin of the exceptional fluorescence properties of this sequence, according to quantum chemical calculations (24,30). Since 5′–5′ stacking of G-tetrads is a basic structural feature for left-handed G4 structures, we may expect that other sequences with this structural feature also display enhanced emission properties.

**Figure 10.**
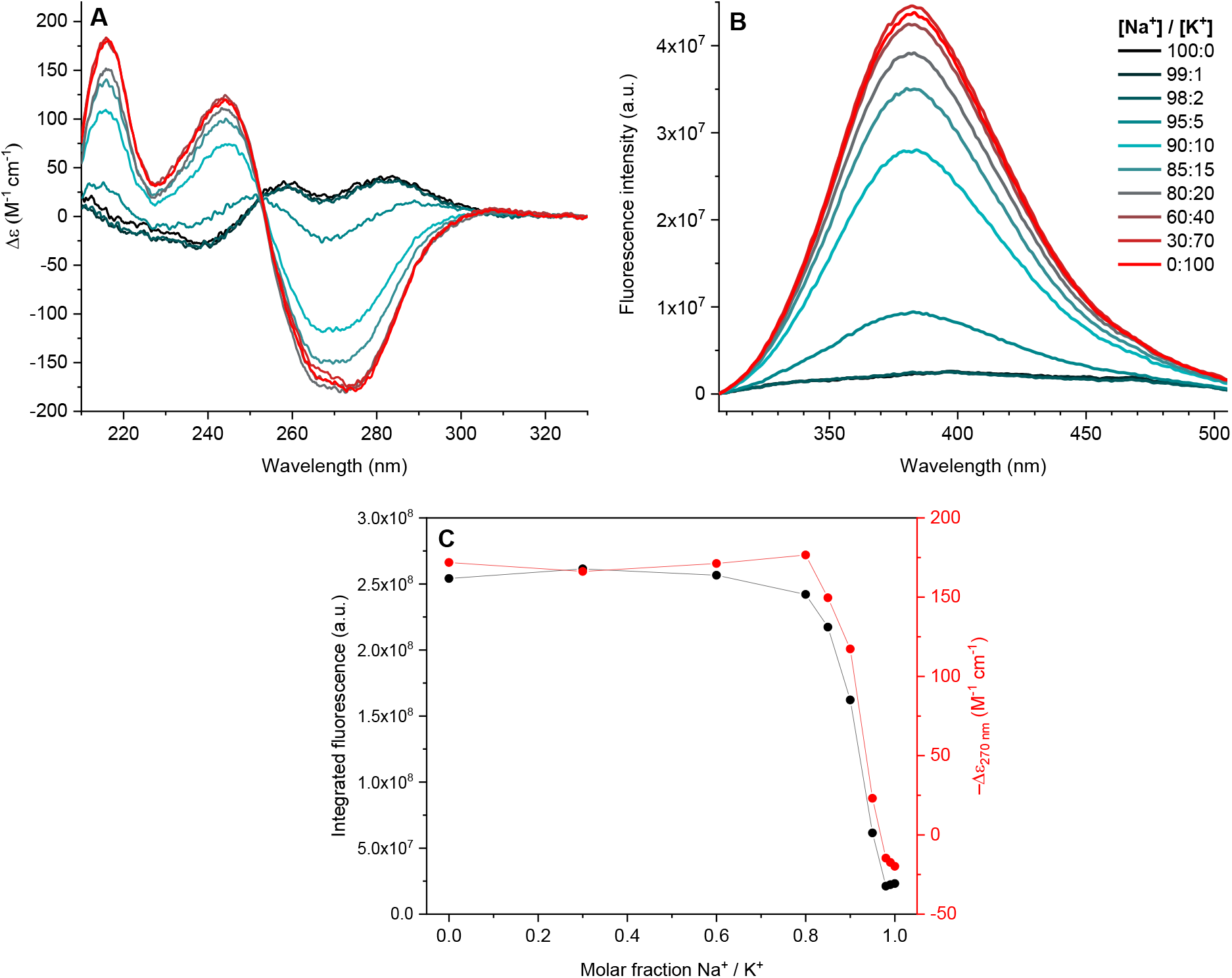
Following the conformational transition of *ZG4* from unfolded state to a left-handed G4 by CD and intrinsic fluorescence. A) CD spectra of *ZG4* solutions (*c* = 5.7 μM) containing a variable proportion of Na^+^ and K^+^ (total NaCl + KCl concentration of 0.1 M, lithium cacodylate buffer 0.01 M, pH 7.2); B) corresponding emission spectra (λ_ex_ = 260 nm); C) comparison of the integrated fluorescence intensity (λ_ex_ = 260 nm, integration between 305 and 505 nm) and molar dichroic absorption at 270 nm (from CD spectra) at various Na^+^ / K^+^ proportions.

**Table 2.** Absolute and relative (with respect to *G*_*3*_*T*) fluorescence quantum yields of selected oligonucleotides.^a^

| Sequence | Buffer | $\Phi \times 10^2$ | $\Phi / \Phi_{G_3T}$ |
| --- | --- | --- | --- |
| 22AG | A | 0.069 | 0.19 |
| 26TTA | A | 0.129 | 0.35 |
| 23TAG | A | 0.076 | 0.20 |
| Bom17 | A | 0.059 | 0.16 |
| TBA | A | 0.078 | 0.21 |
| G4CT | A | 0.150 | 0.40 |
| Pu24T | A | 0.214 | 0.57 |
| 26Ceb | A | 0.040 | 0.11 |
| KRAS-22RT | A | 0.237 | 0.64 |
| ds26 | A | 0.028 | 0.08 |
| hairpin-3 | A | 0.063 | 0.17 |
| ss6 | A | 0.045 | 0.12 |
| RND3 | A | 0.039 | 0.10 |
| EPBC | B | 0.137 | 0.37 |
| i-HRAS2 | B | 0.026 | 0.07 |
| ZG4 | A | 0.195 | 0.52 |
| Block2Δ | A | 0.311 | 0.83 |
| 2xBlock2 | A | 0.270 | 0.73 |
| $G_3T$ | A | 0.372 | 1 |

<sup>a</sup> Conditions: $\lambda_{\text{ex}} = 265$ nm, quantum yield standard: quinine sulfate in 0.5 M $\text{H}_2\text{SO}_4$ ( $\Phi = 0.546$ ).

## DISCUSSION

While the fluorescence of DNA oligonucleotides has been documented since more than a decade, this phenomenon has barely been exploited for structural characterization of secondary structures (31). In this work, we attempted to systematically correlate the secondary structure of oligonucleotides with fluorescence properties using a relatively large training set 89 sequences, whose structures had been unambiguously established by independent methods. First, we confirmed the previous observations demonstrating that steady-state fluorescence spectra are sufficiently sensitive to monitor the conformation transitions of oligonucleotides, such as folding or isomerization of G4 structures (21,23), and extended this observation to the folding of iM structures. These observations corroborate the fact that intrinsic fluorescence can be used as a sensitive method for real-time, unbiased monitoring of conformational changes of oligonucleotides in various settings.

Next, we attempted to identify the characteristic spectral signatures of each conformational group of structures. We observed that intramolecular G4 structures were systematically more fluorescent (on average, 2.5-fold) than other types of structures, in spite of a strong intra-group variance, and display a characteristic shape of fluorescence spectra, featuring a broad, unstructured red edge devoid of additional shoulders (cf. Figure 5). On the other hand, iM structures are characterized by an enhanced fluorescence observed upon 300-nm excitation, which appears in the conditions appropriate for their folding. Further information could be obtained from the multivariate analysis of fluorescence spectra. The data obtained in a single buffer allowed a good discrimination of G4 structures, but were insufficient for the discrimination of iMs, single- and double-stranded structures. However, a combination of the data obtained in two conditions (i.e., favorable and unfavorable for iM folding) allowed a separation of iM-forming sequences as a separate cluster in a PCA plot (cf. Figure 7). Finally, supervised LDA of the reduced dataset (consisting of most informative wavelength readings) allowed a relatively good discrimination (error rate of 9.1%) between the three distinct groups of G4, iM, and other (single- and double-stranded) oligonucleotide conformations (cf. Figure 8). Remarkably, the discrimination between the latter was not possible, neither by naked-eye inspection of the intensity or the shape of their fluorescence spectra (cf. Figures 4 and 5), nor by multivariate analysis of the latter. Notably, CD spectroscopy is not performing better in this case, since the CD spectra of these structural groups are relatively similar; in addition, single-stranded sequences demonstrate a significant intra-group variance in terms of CD spectra (Figure S7, E–F).

The power of supervised multivariate analysis lies in the use of the data obtained with the training set of substrates for the in-class assignment (“typing”) of novel analytes. To test the practical applicability of this approach, we employed four newly generated G-rich sequences and demonstrated that structural predictions based on the LDA of their fluorescence spectra qualitatively agree with the information by two other biophysical techniques, CD and NMR spectroscopy. Furthermore, we assessed several “problematic” sequences adopting unusual secondary structures not included into the training set. In this case, we observed that, in most cases, multivariate analysis of their fluorescent properties correctly assigned these analytes to the group of “other” structures, with a notable exception of non-G4 tetraplex structures (*VK1*, *VK2*, *VK34*), mis-assigned to the G4 group. This fact implies that the fluorescence properties of this class of structures resemble those of G4s, most likely due to similar stacking of guanine residues, and also calls for a detailed investigation of this phenomenon. Notably, three left-handed G4 structures as well as the stacked G4 dimer *G*_*3*_*T* clustered separately but were correctly assigned to the G4 group. This observation implies that the exceptional fluorescent properties of these structures are related to their shared structural feature, that is, the 5′–5′ stacking mode of G-tetrads.

Compared with CD spectroscopy, which is a well-established method for assessment of secondary structures of nucleic acids, the discriminatory power and utility of intrinsic fluorescence may seem more limited at a first glance. Indeed, we were not able to discriminate between different topological classes of G4 structures, which can be easily achieved through the multivariate (16,17) or naked-eye analysis of CD spectra (14). Moreover, the discrimination between the classes of G4s, iMs, and other structures was achieved with a non-negligible error rate of 9.1%. However, we should stress out that discrimination of G4 and iM structures from single-stranded and/or duplex structures is not always trivial with CD spectroscopy (16), and often requires additional methods such as NMR spectroscopy, mass-spectrometry, and/or thermal methods (43). Finally, the key advantage of fluorescence-based methods is the possibility of miniaturization and high-throughput screening, which are not accessible with other methods. Indeed, the implementation of this method in a DNA microarray format will allow the screening and chemometric analysis of up to hundreds of thousands of oligonucleotides in one experiment, paving a way to structural profiling of whole genomes.

## ACKNOWLEDGEMENTS

The authors thank Dr. Jean-Louis Mergny for helpful discussions.

## SUPPLEMENTARY DATA

Supplementary Data are available at NAR online.

