## Supplementary Information for "Harnessing intrinsic fluorescence for typing of secondary structures of DNA"

### CONTENTS


| | |
|---|---|
| **Table S1.** Training set of DNA sequences | S2 |
| **Figure S1.** Comparison of CD and fluorescence emission spectra of folded and unfolded G4-forming sequences | S4 |
| **Figure S2.** Following the folding of *22AG* by CD and intrinsic fluorescence | S5 |
| **Figure S3.** Comparison of integrated fluorescence intensity ($\lambda_{ex}$ = 300 nm) and molar dichroic absorption upon conformational change of *22AG* in solutions with varying $Na^+$ / $K^+$ content | S5 |
| **Figure S4.** Comparison of CD and fluorescence emission spectra of folded and unfolded iM-forming sequences | S6 |
| **Figure S5.** Following the unfolding of *EBPC* iM structure by CD and intrinsic fluorescence | S7 |
| **Figure S6.** CD spectra of tested oligonucleotides: G4-forming sequences | S8 |
| **Figure S7.** CD spectra of tested oligonucleotides: iM-forming sequences, duplexes and single-strand | S9 |
| **Table S2.** Conformational characterization of G4- and iM-forming sequences based on the qualitative inspection of CD spectra in different conditions | S10 |
| **Figure S8.** Emission spectra ($\lambda_{ex}$ = 260 nm) of the tested DNA sequences, grouped according to their expected conformation | S13 |
| **Figure S9.** Emission spectra ($\lambda_{ex}$ = 300 nm) of the tested DNA sequences, grouped according to their expected conformation | S15 |
| **Figure S10.** Group-averaged, normalized emission spectra ($\lambda_{ex}$ = 260 nm, buffer B) of the tested oligonucleotides, grouped according to their expected conformation | S16 |
| **Figure S11.** Group-averaged, normalized emission spectra ($\lambda_{ex}$ = 300 nm, buffers A and B) of the tested oligonucleotides, grouped according to their expected conformation | S17 |
| **Figure S12.** PC1 vs. PC2 plots resulting from the PCA of normalized emission spectra recorded in buffer A ($\lambda_{ex}$ = 300 nm) and in buffer B ($\lambda_{ex}$ = 300 nm) | S18 |
| **Figure S13.** Scree plot of the eigenvalues obtained for the PCA shown in Figure 6, A | S18 |
| **Figure S14.** Analysis of the variance of the emission dataset | S19 |
| **Figure S15.** 3D PCA plots resulting from the analysis of reduced datasets | S19 |
| **Table S3.** Sequences and LDA-based conformational assignment of test sequences *RND-HS1* to *RND-HS4* | S20 |
| **Figure S16.** 3D PCA and LDA plots obtained from the analysis of the training set and test sequences *RND-HS1* to *RND-HS4* | S20 |
| **Figure S17.** $^1$H NMR spectra of oligonucleotides *RND-HS1* to *RND-HS4* | S20 |
| **Figure S18.** CD and TDS spectra of test sequences *RND-HS1* to *RND-HS4* | S21 |
| **Figure S19.** CD spectra of oligonucleotides adopting peculiar conformations in buffers A and B | S21 |
| **Table S4.** LDA-based assignment of the test sequences presented in Figure S18 | S22 |
| **Figure S20.** Fluorescence emission spectra of left-handed G4s | S22 |
| **Figure S21.** Determination of fluorescence quantum yields | S25 |
| **Figure S22.** Structural details of a typical parallel G4 (*c-myc*), left-handed G4 (*ZG4*) and a stacked parallel dimeric G4 (*J19*, an analogue of $G_3T$) | S26 |

**Table S1.** Training set of DNA sequences.

| Acronym | Sequence (5′→ 3′) | Reported conformation | PDB entry |
|---|---|---|---|
| *22AG* | AGGGTTAGGGTTAGGGTTAGGG | hybrid G4 | - |
| *46AG* | AGGGTTAGGGTTAGGGTTAGGGTTAGGGTTAGGG TTAGGGTTAGGG | hybrid G4 | - |
| *Bcl2Mid* | GGGCGCGGGAGGAATTGGGCGGG | hybrid G4 | 2F8U |
| *UpsB-Q3* | CAGGGTTAAGGGTATACATTTAGGGGTTAGGGTT | hybrid G4 | - |
| *26TTA* | TTAGGGTTAGGGTTAGGGTTAGGGTT | hybrid G4 | 2JPZ |
| *25TAG* | TAGGGTTAGGGTTAGGGTTAGGGTT | hybrid G4 | 2JSL |
| *23TAG* | TAGGGTTAGGGTTAGGGTTAGGG | hybrid G4 | 2JSK |
| *24TTA* | TTAGGGTTAGGGTTAGGGTTAGGG | hybrid G4 | 2JSL |
| *VEGFR-17T* | GGGTACCCGGGTGAGGTGCGGGGT | hybrid G4 | 5ZEV |
| *TP3-T6* | TGGGGTCCGAGGCGGGGCTTGGG | hybrid G4 | 6AC7 |
| *chl1* | GGGTGGGGAAGGGGTGGGT | hybrid G4 | 2KPR |
| *UpsB-Q1* | CAGGGTTAAGGGTATAACTTTAGGGGTTAGGGTT | hybrid G4 | 5MTA |
| *LTR-III* | GGGAGGCGTGGCCTGGGCGGGACTGGGG | hybrid G4 | 6H1K |
| *SP-PGQ-1* | GGGCAACTTGGCTGGGGTCTAGTTCCACGGGACG GG | hybrid G4 | - |
| *26CEB* | AAGGGTGGGTGTAAGTGTGGGTGGGT | parallel G4 | 2LPW |
| *c-kit2-T12T21* | CGGGCGGGCGCTAGGGAGGGT | parallel G4 | 2KYP |
| *KRAS-22RT* | AGGGCGGTGTGGGAATAGGGAA | parallel G4 | 5I2V |
| *Pu24T* | TGAGGGTGGTGAGGGTGGGGAAGG | parallel G4 | 2A5P |
| *c-kit87up* | AGGGAGGGCGCTGGGAGGAGGG | parallel G4 | 2O3M |
| *VEGF* | CGGGGCGGGCCTTGGGCGGGGT | parallel G4 | 2M27 |
| *c-myc* | TGAGGGTGGGTAGGGTGGGTAA | parallel G4 | 1XAV |
| *T95-2T* | TTGGGTGGGTGGGTGGGT | parallel G4 | 2LK7 |
| *SP-PGQ-2* | GGGCTAGTGGGGGGAGGGGG | parallel G4 | - |
| *SP-PGQ-3* | GGGCTAATAGGGAGAGCAGGGACGGGG | parallel G4 | - |
| *PCNA G4* | CAGGGCGACGGGGGCGGGGCGGGGCG | parallel G4 | - |
| *TB-1* | TTGTGGTGGGTGGGTGGGT | parallel G4 | 2M4P |
| *T2B-1* | TTGTTGGTGGGTGGGTGGGT | parallel G4 | - |
| *TB-3* | TTGGGTGTGGTGGGTGGGT | parallel G4 | - |
| *G15* | TTGGGGGGGGGGGGGGGT | parallel G4 | 2MB2 |
| *Myc1245* | TTGGGGAGGGTTTTAAGGGTGGGGAAT | parallel G4 | 6NEB |
| *AT11* | TGGTGGTGGTTGTTGTGGTGGTGGTGGT | parallel G4 | 2N3M |
| *LTR-IV* | CTGGGCGGGACTGGGGAGTGGT | parallel G4 | 2N4Y |
| *hras-1* | TCGGGTTGCGGGCGCAGGGCACGGGCG | anti-parallel G4 | - |
| *TBA* | GGTTGGTGTGGTTGG | anti-parallel G4 | 148D |
| *HIV-PRO-1* | TGGCCTGGGCGGGACTGGG | anti-parallel G4 | - |
| *22CTA* | AGGGCTAGGGCTAGGGCTAGGG | anti-parallel G4 | - |
| *Bm-U16* | TAGGTTAGGTTAGGTUAGG | anti-parallel G4 | - |
| *c-kit** | GGCGAGGAGGGGCGTGGCCGGC | anti-parallel G4 | 6GH0 |
| *Bom17* | GGTTAGGTTAGGTTAGG | anti-parallel G4 | - |
| *G4CT* | GGGGCTGGGGCTGGGGCTGGGG | anti-parallel G4 | - |
| *22GGG* | GGGTTAGGGTTAGGGTTAGGGT | anti-parallel G4 | 2KF8 |
| *htel21T18* | GGGTTAGGGTTAGGGTTTGGG | anti-parallel G4 | 5YEY |
| *19wt* | GGGGGAGGGGTACAGGGGTACAGGGG | anti-parallel G4 | 6FTU |
| *LWDLN 1* | GGGTTTGGGTTTTGGGAGGG | anti-parallel G4 | 5J05 |
| *LWDLN 2* | GGGGTTGGGGTTTTGGGGAAGGGG | anti-parallel G4 | 2M6W |
| *LWDLN 3* | GGTTTGGTTTTGGTTGG | anti-parallel G4 | 5J4W |
| *ss 1* | CACTAAACCTAACACTAACCAT | single strand | - |
| *ss 9* | ATGGTTAGTGTTAGGTTTAGTG | single strand | - |
| *ss 3* | GTCGCCGGGCCAGTCGTCCATAC | single strand | - |
| *ss 4* | GTATGGACGACTGGCCCGGCGAC | single strand | - |

| | | | |
|---|---|---|---|
| *ss 6* | GACGTGTCGAAAGAGCTCCGATTA | single strand | - |
| *ss 7* | TAATCGGAGCTCTTTCGACACGTC | single strand | - |
| *RND1* | CTATACGAAAACCTTTTGTATCATT | single strand | - |
| *RND2* | AATGATACAAAAGGTTTTCGTATAG | single strand | - |
| *RND3* | TAACGTTTATAATGTAGTCTCATTA | single strand | - |
| *RND4* | TAATGAGACTACATTATAAACGTTA | single strand | - |
| *RND5* | AGAATTATTCGGGGGCAATGACAAC | single strand | - |
| *RND6* | GTTGTCATTGCCCCCGAATAATTCT | single strand | - |
| *RND7* | GCCTTGCGGAGGCATGCGTCATGCT | single strand | - |
| *RND8* | AGCATGACGCATGCCTCCGCAAGGC | single strand | - |
| *dT26* | TTTTTTTTTTTTTTTTTTTTTTTTTT | single strand | - |
| *ctDNA* | genomic, highly polymerized | duplex | - |
| *ds26* | CAATCGGATCGAATTCGATCCGATTG | duplex | - |
| *ds-lac* | GAATTGTGAGCGCTCACAATTC | duplex | - |
| *Hairpin 1* | GGATTCTTGGATTTTCCAAGAATCC | duplex | - |
| *duplex 19* | CACTAAACCTAACACTAACCAT<br>ATGGTTAGTGTTAGGTTTAGTG | duplex | - |
| *duplex 34* | GTCGCCGGGCCAGTCGTCCATAC<br>GTATGGACGACTGGCCCGGCGAC | duplex | - |
| *duplex 67* | GACGTGTCGAAAGAGCTCCGATTA<br>TAATCGGAGCTCTTTCGACACGTC | duplex | - |
| *RND1+2* | CTATACGAAAACCTTTTGTATCATT<br>AATGATACAAAAGGTTTTCGTATAG | duplex | - |
| *RND3+4* | TAACGTTTATAATGTAGTCTCATTA<br>TAATGAGACTACATTATAAACGTTA | duplex | - |
| *Hairpin 2* | TCGGTATTGTGTTTCACAATACCGA | duplex | - |
| *RND7+8* | AGAATTATTCGGGGGCAATGACAAC<br>GTTGTCATTGCCCCCGAATAATTCT | duplex | - |
| *RND5+6* | GCCTTGCGGAGGCATGCGTCATGCT<br>AGCATGACGCATGCCTCCGCAAGGC | duplex | - |
| *Hairpin 3* | AGGACGGTGTATTTTACACCGTCCT | duplex | - |
| *DDD* [a] | CGCGAATTCGCG | duplex | 2BNA |
| *i-hTel* | CCCTAACCCTAACCCTAACCCT | iM | - |
| *21rHTS* | CCCAATCCCAATCCCAATCCC | iM | - |
| *i-HRAS1* | CGCCCGTGCCCTGCGCCCGCAACCCGA | iM | - |
| *i-HRAS2* | ACCGCGCGCCCCCGCCCCCGCCCCGCCCCGGCC<br>TCG | iM | - |
| *i-bcl2* | CAGCCCGCTCCCGCCCCCTTCCTCCCGCGCCCGC<br>CCCT | iM | - |
| *EBPC* | CCCCTCCCCTCCCCTCCCC | iM | - |
| *OBPC* | CCCCTCCCTTTCCCTCCCC | iM | - |
| *Py27 (1245)* | AATCCCCTCCCATTTTTCCCACCCCTT | iM | - |
| *Py22(2345)* | AATCCCACCCCTCCCACCCCTT | iM | - |
| *dC20* | TCCCCCCCCCCCCCCCCCCC | iM | - |
| *C3T3* | CCCTTTCCCTTTCCCTTTCCC | iM | - |
| *C4T3* | CCCCTTTCCCCTTTCCCCTTTCCCC | iM | - |
| *C5T3* | CCCCCTTTCCCCCTTTCCCCCTTTCCCCC | iM | - |
| *C3A3* | CCCAAACCCAAACCCAAACCC | iM | - |

[a] Dickerson–Drew dodecamer.

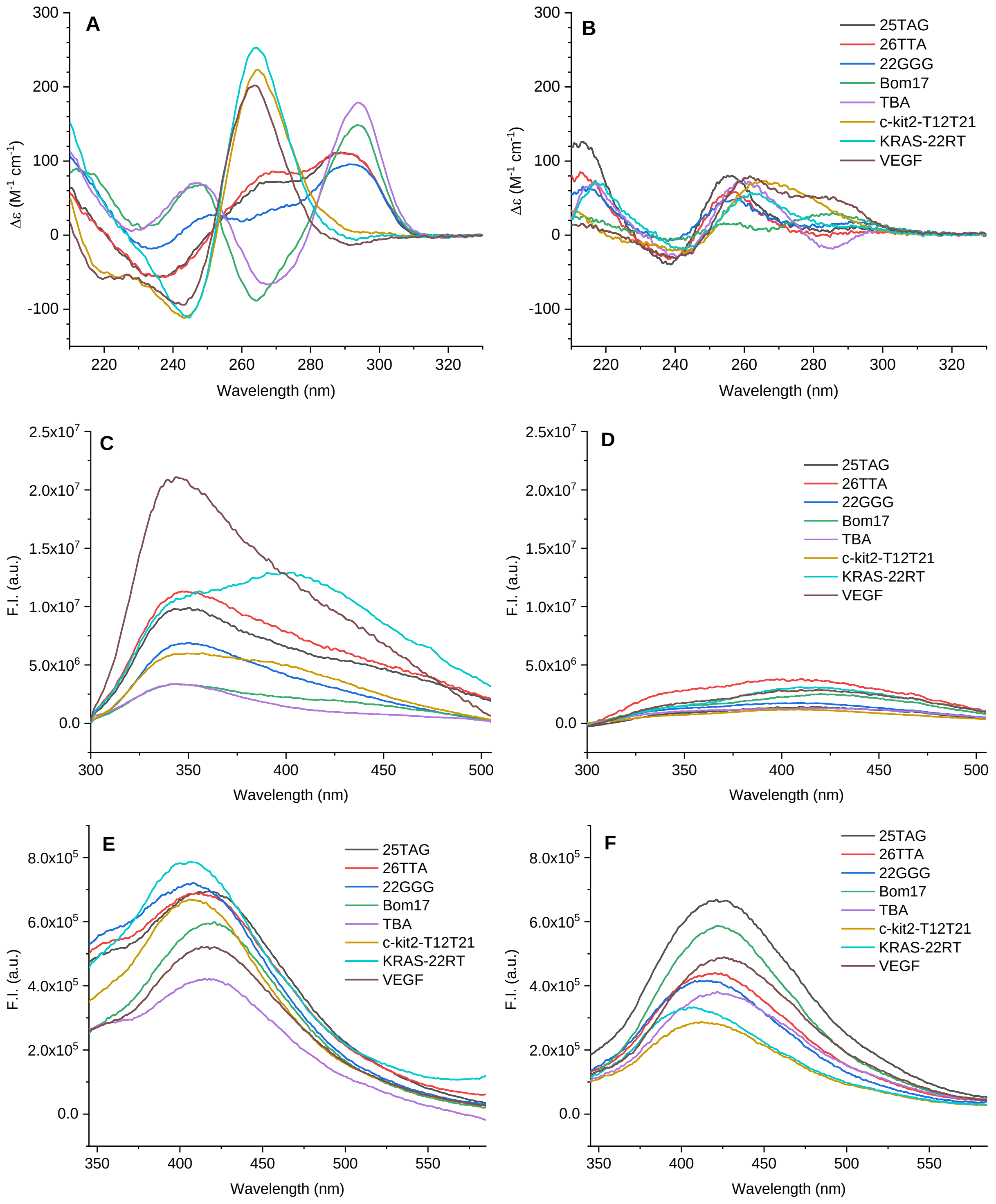


**Figure S1.** Comparison of CD and fluorescence emission spectra of folded and unfolded G4-forming sequences. A) CD spectra of 5.7 µM oligonucleotides in buffer A; B) CD spectra of 5.7 µM oligonucleotides in a $K^+$-free buffer (0.1 M LiCl, 0.01 M lithium cacodylate, pH 7.2); C,E) fluorescence spectra (C: $\lambda_{ex}$ = 260 nm, E: $\lambda_{ex}$ = 300 nm) of samples presented in A; D,F) fluorescence spectra (D: $\lambda_{ex}$ = 260 nm, F: $\lambda_{ex}$ = 300 nm) of the samples presented in B.

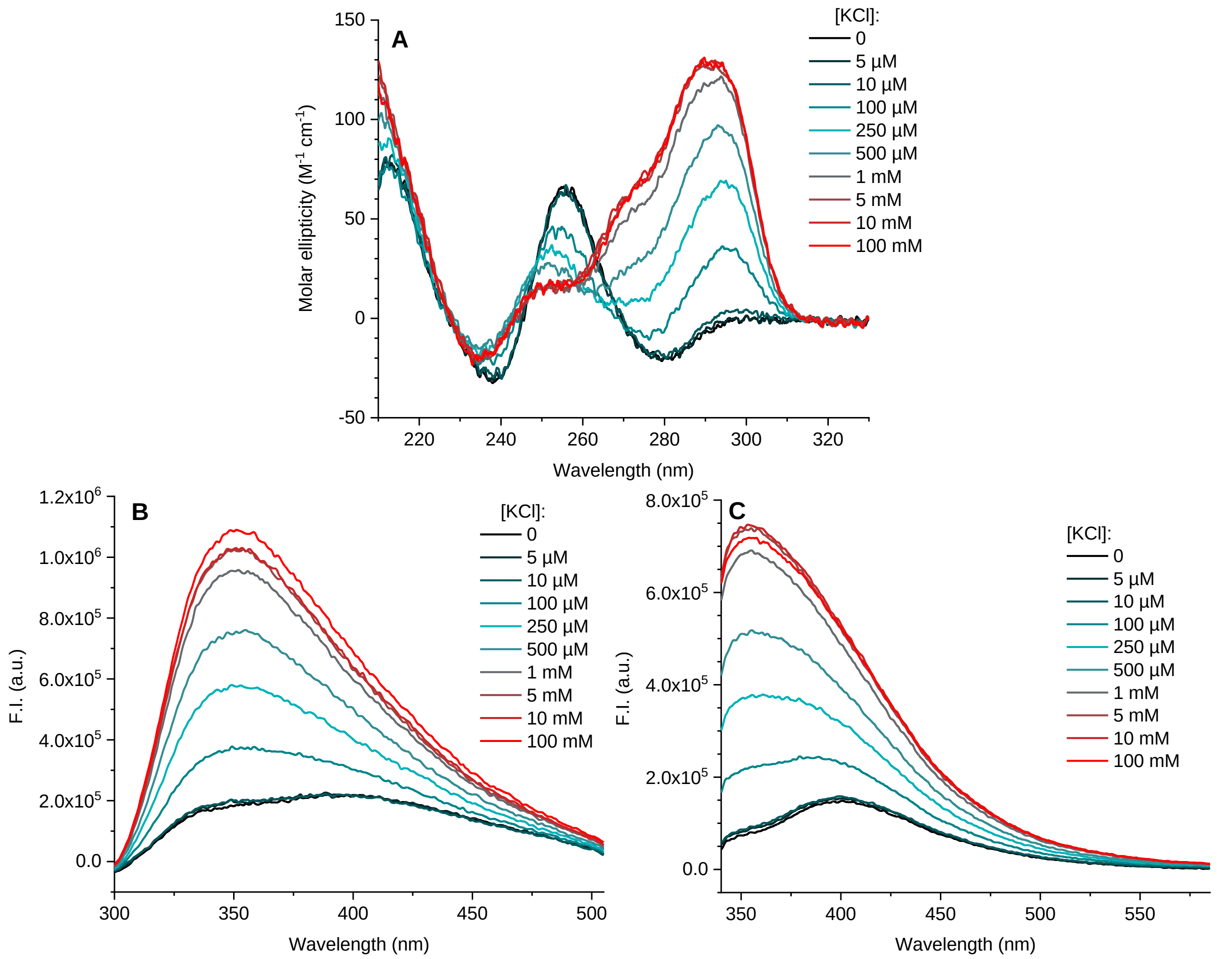


**Figure S2.** Following the folding of *22AG* by CD and intrinsic fluorescence. A) CD spectra of *22AG* ($c$ = 5.7 µM) in buffer solutions (0.01 M lithium cacodylate, pH 7.2) with variable KCl content. B–C) Corresponding fluorescence spectra obtained with B) $\lambda_{ex}$ = 260 nm and C) $\lambda_{ex}$ = 300 nm. Fluorescence spectra were corrected for the inner filter effect.

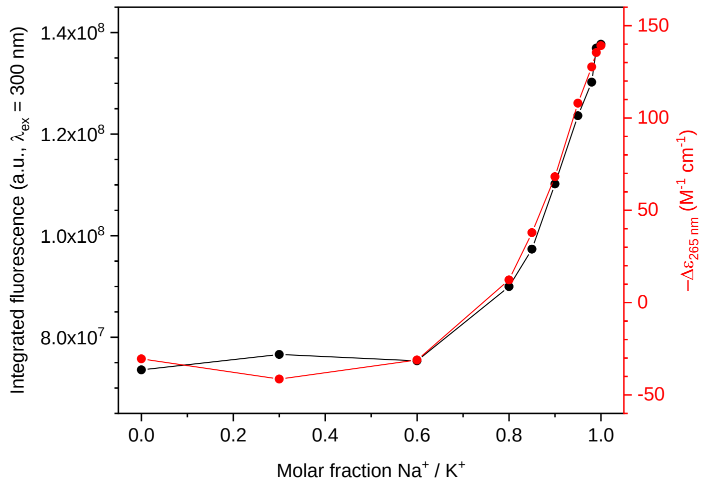


**Figure S3.** Comparison of integrated fluorescence intensity ($\lambda_{ex}$ = 300 nm) and molar dichroic absorption at 265 nm (from CD spectra, red lines and dots) upon conformational change of *22AG* ($c$ = 5.7 µM) in buffer solutions with varying $Na^+$ / $K^+$ content.

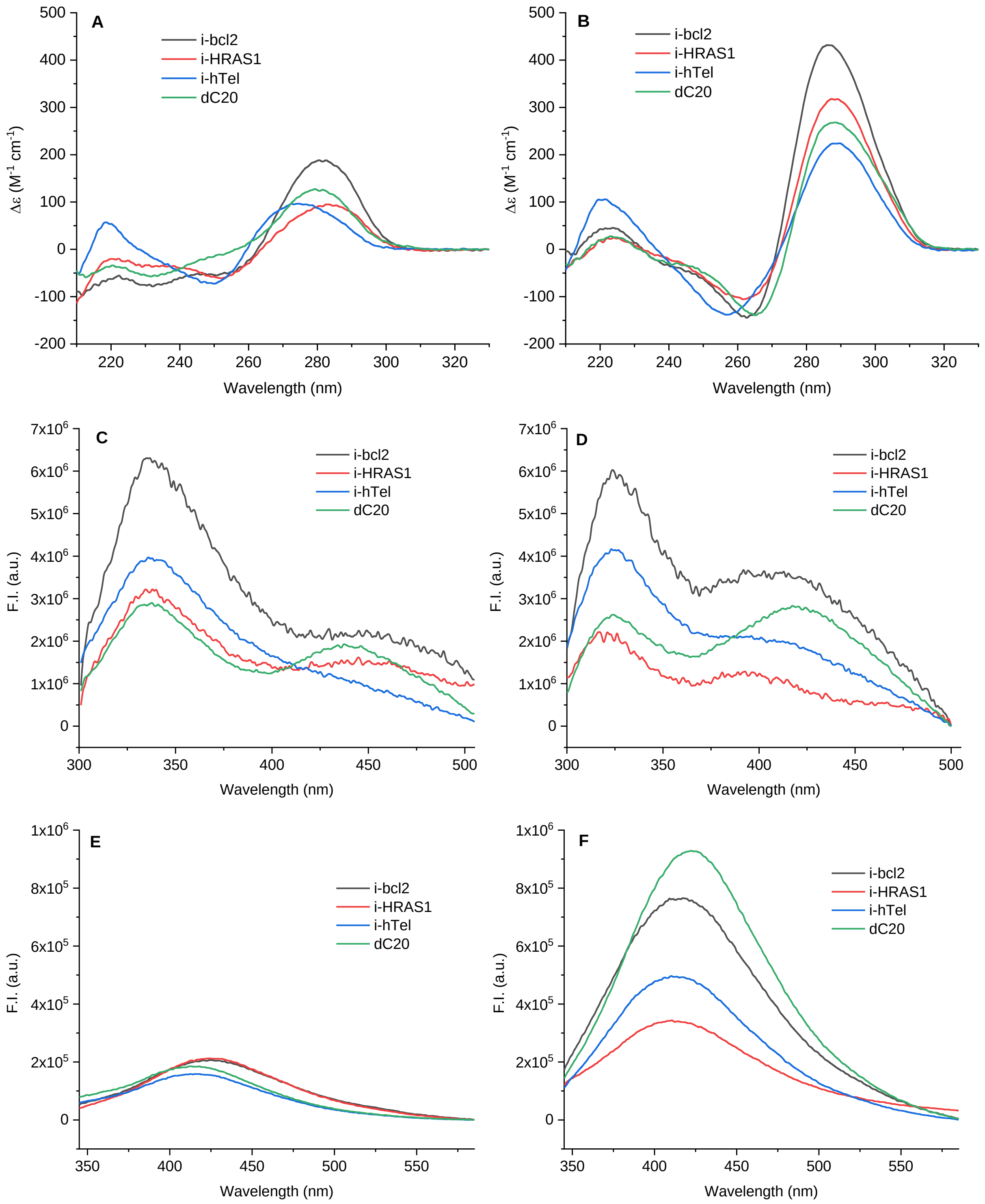


**Figure S4.** Comparison of CD and fluorescence emission spectra of folded and unfolded iM-forming sequences. A) CD spectra of 5.7 µM oligonucleotides in buffer A; B) CD spectra of 5.7 oligonucleotides in buffer B; C,E) fluorescence spectra (C: $\lambda_{ex}$ = 260 nm, E: $\lambda_{ex}$ = 300 nm) of samples presented in A; D,F) fluorescence spectra (D: $\lambda_{ex}$ = 260 nm, F: $\lambda_{ex}$ = 300 nm) of the samples presented in B. Fluorescence spectra were corrected for the inner filter effect.

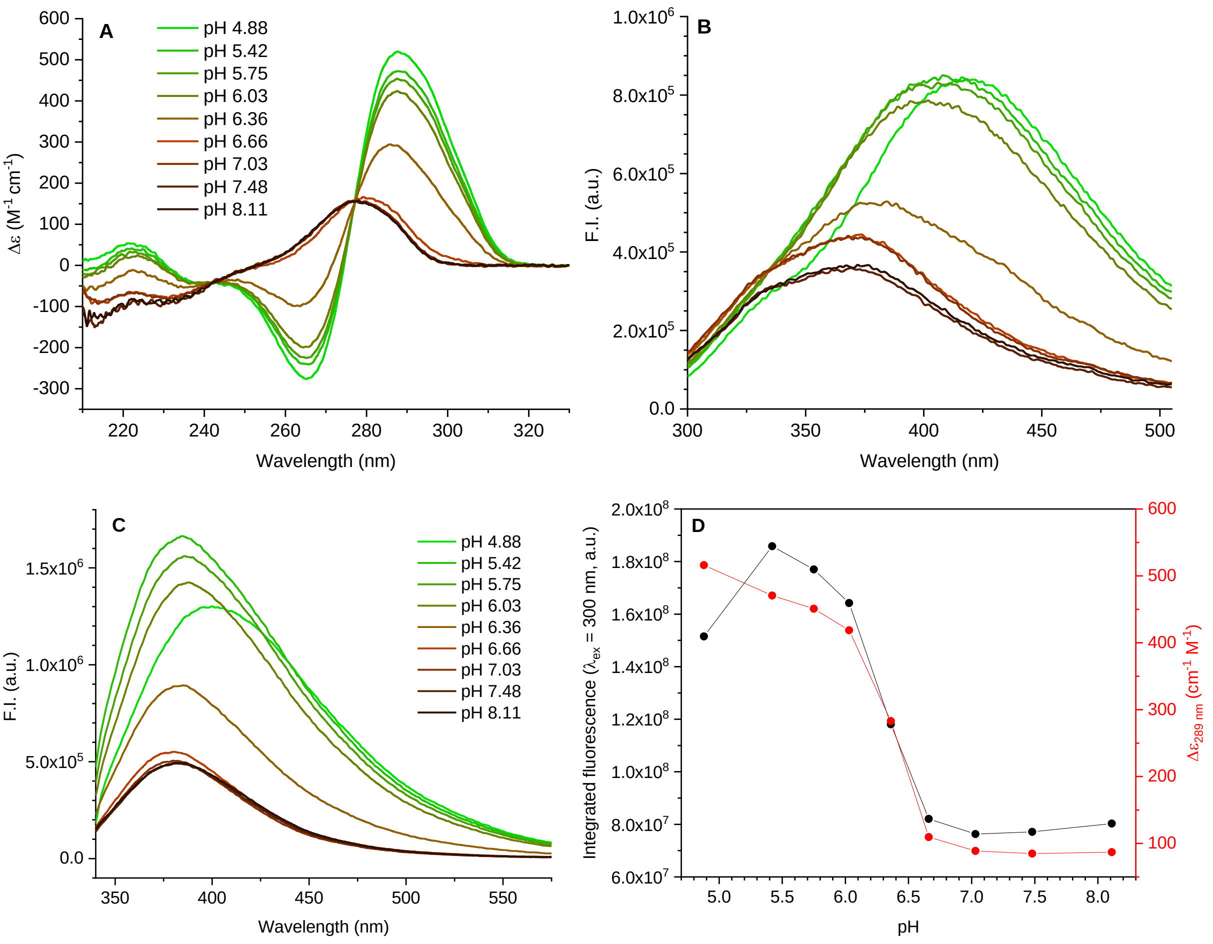


**Figure S5.** Following the unfolding of *EBPC* iM structure by CD and intrinsic fluorescence. A) CD spectra of *EPBC* solutions ($c$ = 5.7 μM) with increasing pH (4.9–8.1, 0.01 lithium cacodylate buffer, 0.1 M KCl); B–C) corresponding fluorescence emission spectra obtained with B) $\lambda_{ex}$ = 260 nm and C) $\lambda_{ex}$ = 300 nm; D) comparison of the integrated fluorescence emission ($\lambda_{ex}$ = 300 nm) and molar dichroic absorption at 289 nm (from CD spectra, red lines and dots) of the *EPBC* samples presented above. Fluorescence spectra were corrected for the inner filter effect.

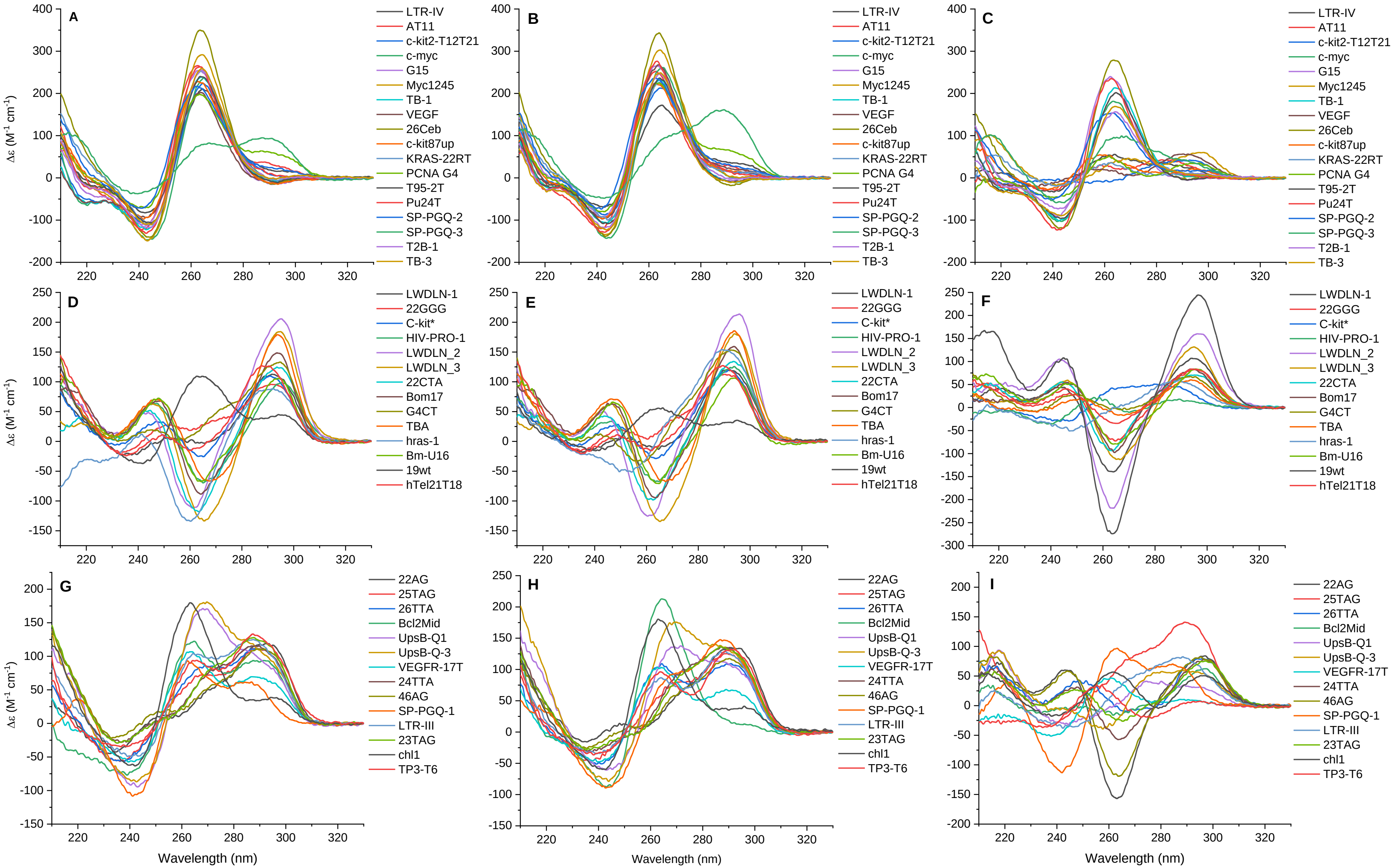


**Figure S6.** CD spectra of the tested oligonucleotides: G4-forming sequences ($c$ = 5.7 µM). A–C) Expected parallel G4 structures in A) buffer A, B) buffer B and C) buffer C; D–F) expected anti-parallel G4 structures in D) buffer A, E) buffer B and F) buffer C; G–I) expected hybrid G4 structures in G) buffer A, H) buffer B and I) buffer C.

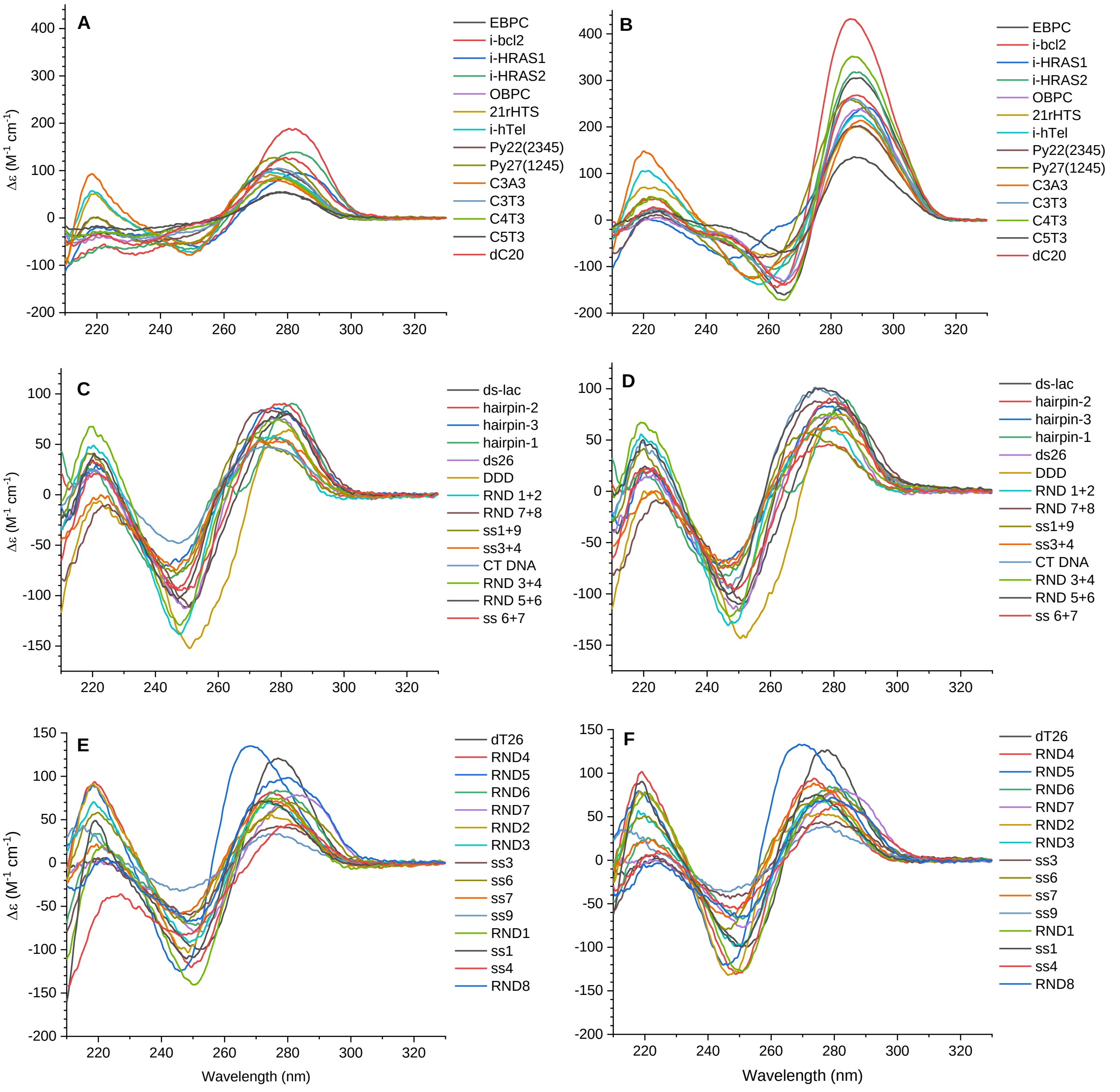


**Figure S7.** CD spectra of the tested oligonucleotides: iM-forming sequences, duplexes and single-strands ($c$ = 5.7 µM). A–B) iM-forming sequences in A) buffer A and B) buffer B; C–D) double-stranded structures in C) buffer A and D) buffer B; E–F) single-stranded sequences in E) buffer A and F) buffer B.

**Table S2.** Conformational characterization of G4- and iM-forming sequences based on the qualitative inspection of CD spectra in different conditions.

| **Acronym** | **Reported structure** [a] | **Buffer A** | **Buffer B** | **Buffer C** |
|---|---|---|---|---|
| 22AG | hybrid G4 | hybrid G4 | hybrid G4 | anti-parallel G4 |
| 46AG | hybrid G4 | hybrid G4 | hybrid G4 | anti-parallel G4 |
| Bcl2Mid | hybrid G4 | hybrid G4 | parallel G4 | [d] |
| UpsB-Q-3 | hybrid G4 | hybrid G4 | hybrid G4 | anti-parallel G4 [d] |
| 26TTA | hybrid G4 | hybrid G4 | hybrid G4 | anti-parallel G4 [d] |
| 25TAG | hybrid G4 | hybrid G4 | hybrid G4 | hybrid G4 |
| 23TAG | hybrid G4 | hybrid G4 | hybrid G4 | anti-parallel G4 [d] |
| 24TTA | hybrid G4 | hybrid G4 | hybrid G4 | anti-parallel G4 |
| VEGFR-17T | hybrid G4 | hybrid G4 | hybrid G4 | partially unfolded |
| TP3-T6 | hybrid G4 | hybrid G4 | hybrid G4 | [d] |
| chl1 | hybrid G4 | hybrid G4 | hybrid G4 | anti-parallel G4 [d] |
| UpsB-Q1 | hybrid G4 | hybrid G4 | hybrid G4 | unfolded [d] |
| LTR-III | hybrid G4 | hybrid G4 | hybrid G4 | partially unfolded |
| SP-PGQ-1 | hybrid G4 | hybrid G4 | hybrid G4 [d] | hybrid G4 |
| 26CEB | parallel G4 | parallel G4 | parallel G4 | parallel G4 |
| c-kit2-T12T21 | parallel G4 | parallel G4 | parallel G4 | anti-parallel G4 [d] |
| KRAS-22RT | parallel G4 | parallel G4 | parallel G4 | unfolded |
| Pu24T | parallel G4 | parallel G4 | parallel G4 | parallel G4 |
| c-kit87up | parallel G4 | parallel G4 | parallel G4 | unfolded [d] |
| VEGF | parallel G4 | parallel G4 | parallel G4 | anti-parallel G4 [d] |
| c-myc | parallel G4 | parallel G4 | parallel G4 | parallel G4 |
| T95-2T | parallel G4 | parallel G4 | parallel G4 | parallel G4 |
| SP-PGQ-2 | parallel G4 | parallel G4 | parallel G4 | partially unfolded |
| SP-PGQ-3 | parallel G4 | hybrid G4 | hybrid G4 [d] | parallel G4 [d] |
| PCNA G4 | parallel G4 | parallel G4 [d] | parallel G4 [d] | unfolded [d] |
| TB-1 | parallel G4 | parallel G4 | parallel G4 | parallel G4 |
| T2B-1 | parallel G4 | parallel G4 | parallel G4 | partially unfolded |
| TB-3 | parallel G4 | parallel G4 | parallel G4 | partially unfolded |
| G15 | parallel G4 | parallel G4 | parallel G4 | parallel G4 |
| Myc1245 | parallel G4 | parallel G4 | parallel G4 | anti-parallel G4 [d] |
| AT11 | parallel G4 | parallel G4 | parallel G4 | unfolded |
| LTR-IV | parallel G4 | parallel G4 | parallel G4 | unfolded |
| hras-1 | anti-parallel G4 | anti-parallel G4 | [d] | [d] |
| TBA | anti-parallel G4 | anti-parallel G4 | anti-parallel G4 | [d] |
| HIV-PRO-1 | anti-parallel G4 | anti-parallel G4 | anti-parallel G4 | [d] |
| 22CTA | anti-parallel G4 | anti-parallel G4 | anti-parallel G4 | anti-parallel G4 |
| Bm-U16 | anti-parallel G4 | anti-parallel G4 | anti-parallel G4 | anti-parallel G4 |
| c-kit* | anti-parallel G4 | anti-parallel G4 | anti-parallel G4 | [d] |
| Bom17 | anti-parallel G4 | anti-parallel G4 | anti-parallel G4 | anti-parallel G4 |
| G4CT | anti-parallel G4 | anti-parallel G4 | anti-parallel G4 | partially unfolded |
| 22GGG | anti-parallel G4 | [d] | [d] | anti-parallel G4 |
| htel21T18 | anti-parallel G4 | anti-parallel G4 | anti-parallel G4 | anti-parallel G4 [d] |
| 19wt | anti-parallel G4 [b] | hybrid G4 [d] | hybrid G4 [d] | anti-parallel G4 |
| LWDLN 1 | anti-parallel G4 [c] | [d] | [d] | anti-parallel G4 |
| LWDLN 2 | anti-parallel G4 [c] | anti-parallel G4 | anti-parallel G4 | anti-parallel G4 |
| LWDLN 3 | anti-parallel G4 [c] | anti-parallel G4 | anti-parallel G4 | anti-parallel G4 |
| i-hTel | iM | unfolded | iM | n/a [e] |
| 21rHTS | iM | unfolded | iM | n/a |
| i-HRAS1 | iM | unfolded | iM | n/a |
| i-HRAS2 | iM | unfolded | iM | n/a |
| i-bcl2 | iM | unfolded | iM | n/a |

| | | | | |
|---|---|---|---|---|
| EBPC | iM | unfolded | iM | n/a |
| OBPC | iM | unfolded | iM | n/a |
| Py27 (1245) | iM | unfolded | iM | n/a |
| Py22(2345) | iM | unfolded | iM | n/a |
| dC20 | iM | unfolded | iM | n/a |
| C3T3 | iM | unfolded | iM | n/a |
| C4T3 | iM | unfolded | iM | n/a |
| C5T3 | iM | unfolded | iM | n/a |
| C3A3 | iM | unfolded | iM | n/a |

[a] Unless stated otherwise, the "reported" structures refer to solution structures formed in $K^+$-containing conditions. [b] Crystal structure. [c] In 100 mM $Na^+$, pH 6.8. [d] The CD spectrum could not be interpreted unambiguously or the conformational change was incomplete. [e] Not assessed.

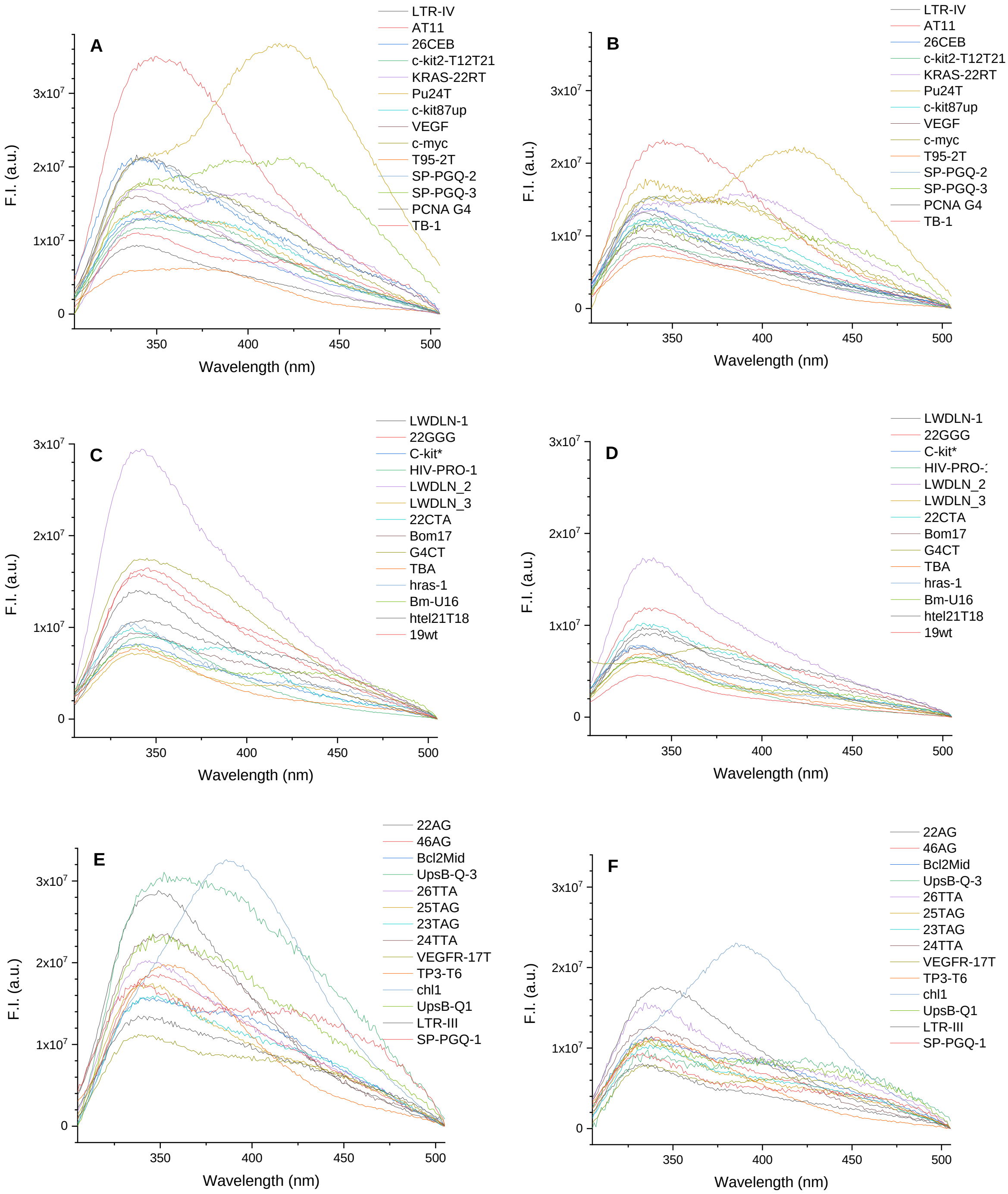
A
B
C
D
E
F
F.I. (a.u.)
Wavelength (nm)
0
1x10^7
2x10^7
3x10^7
350
400
450
500
LTR-IV
AT11
26CEB
c-kit2-T12T21
KRAS-22RT
Pu24T
c-kit87up
VEGF
c-myc
T95-2T
SP-PGQ-2
SP-PGQ-3
PCNA G4
TB-1
LWDLN-1
22GGG
C-kit*
HIV-PRO-1
LWDLN_2
LWDLN_3
22CTA
Bom17
G4CT
TBA
hras-1
Bm-U16
htel21T18
19wt
22AG
46AG
Bcl2Mid
UpsB-Q-3
26TTA
25TAG
23TAG
24TTA
VEGFR-17T
TP3-T6
chl1
UpsB-Q1
LTR-III
SP-PGQ-1

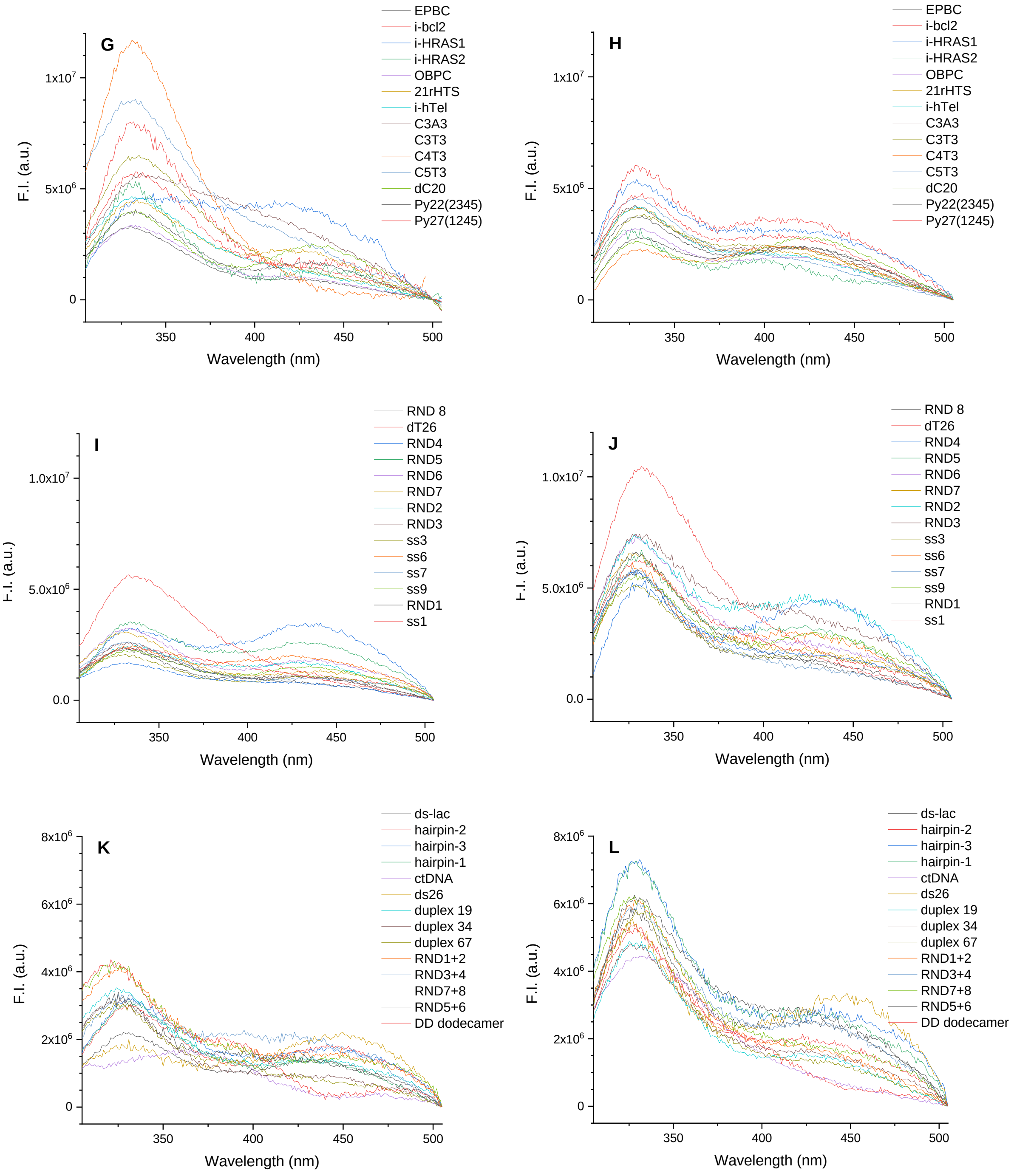


**Figure S8.** Emission spectra ($\lambda_{ex}$ = 260 nm) of the tested DNA sequences ($c$ = 5.7 μM), grouped according to their expected conformation: A–B) parallel G4; C–D) anti-parallel G4; E–F) hybrid G4; G–H) iM; I–J) single strands; K–L) duplex, in buffers A (A, C, E, G, I, K) and B (B, D, F, H, J, L).

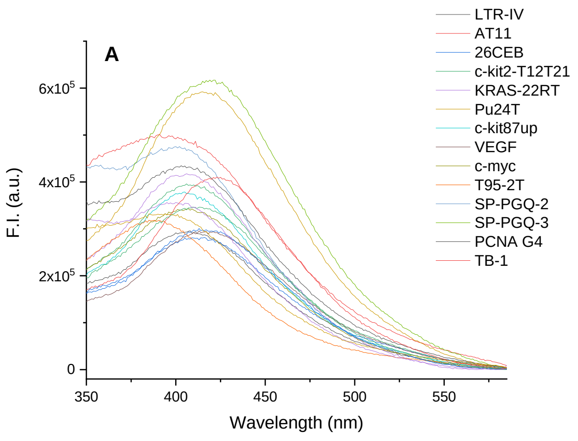
A
F.I. (a.u.)
Wavelength (nm)
LTR-IV
AT11
26CEB
c-kit2-T12T21
KRAS-22RT
Pu24T
c-kit87up
VEGF
c-myc
T95-2T
SP-PGQ-2
SP-PGQ-3
PCNA G4
TB-1

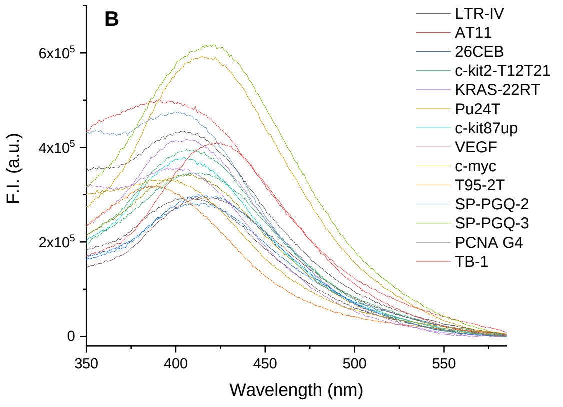
B
F.I. (a.u.)
Wavelength (nm)
LTR-IV
AT11
26CEB
c-kit2-T12T21
KRAS-22RT
Pu24T
c-kit87up
VEGF
c-myc
T95-2T
SP-PGQ-2
SP-PGQ-3
PCNA G4
TB-1

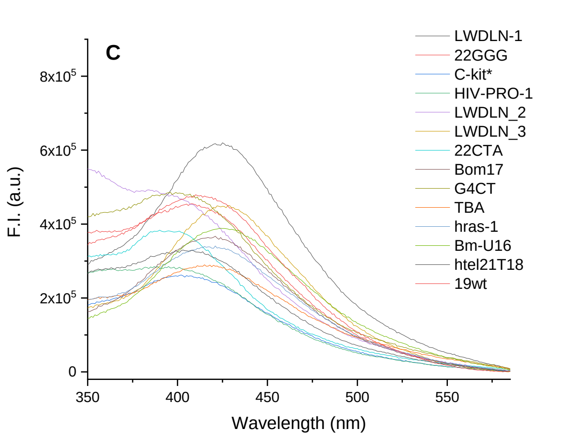
C
F.I. (a.u.)
Wavelength (nm)
LWDLN-1
22GGG
C-kit*
HIV-PRO-1
LWDLN_2
LWDLN_3
22CTA
Bom17
G4CT
TBA
hras-1
Bm-U16
htel21T18
19wt

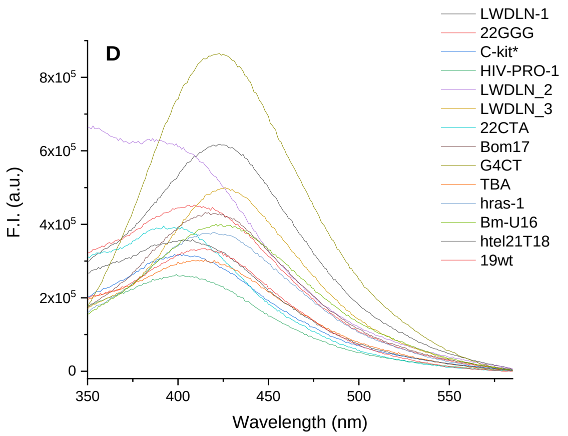
D
F.I. (a.u.)
Wavelength (nm)
LWDLN-1
22GGG
C-kit*
HIV-PRO-1
LWDLN_2
LWDLN_3
22CTA
Bom17
G4CT
TBA
hras-1
Bm-U16
htel21T18
19wt

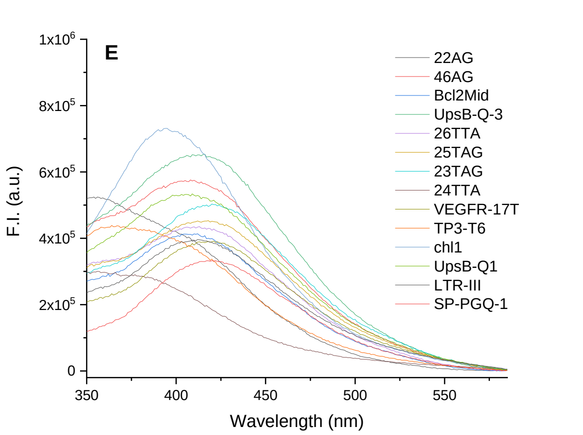
E
F.I. (a.u.)
Wavelength (nm)
22AG
46AG
Bcl2Mid
UpsB-Q-3
26TTA
25TAG
23TAG
24TTA
VEGFR-17T
TP3-T6
chl1
UpsB-Q1
LTR-III
SP-PGQ-1

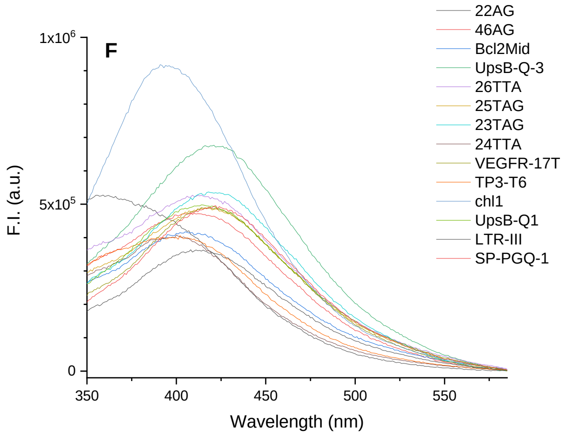
F
F.I. (a.u.)
Wavelength (nm)
22AG
46AG
Bcl2Mid
UpsB-Q-3
26TTA
25TAG
23TAG
24TTA
VEGFR-17T
TP3-T6
chl1
UpsB-Q1
LTR-III
SP-PGQ-1

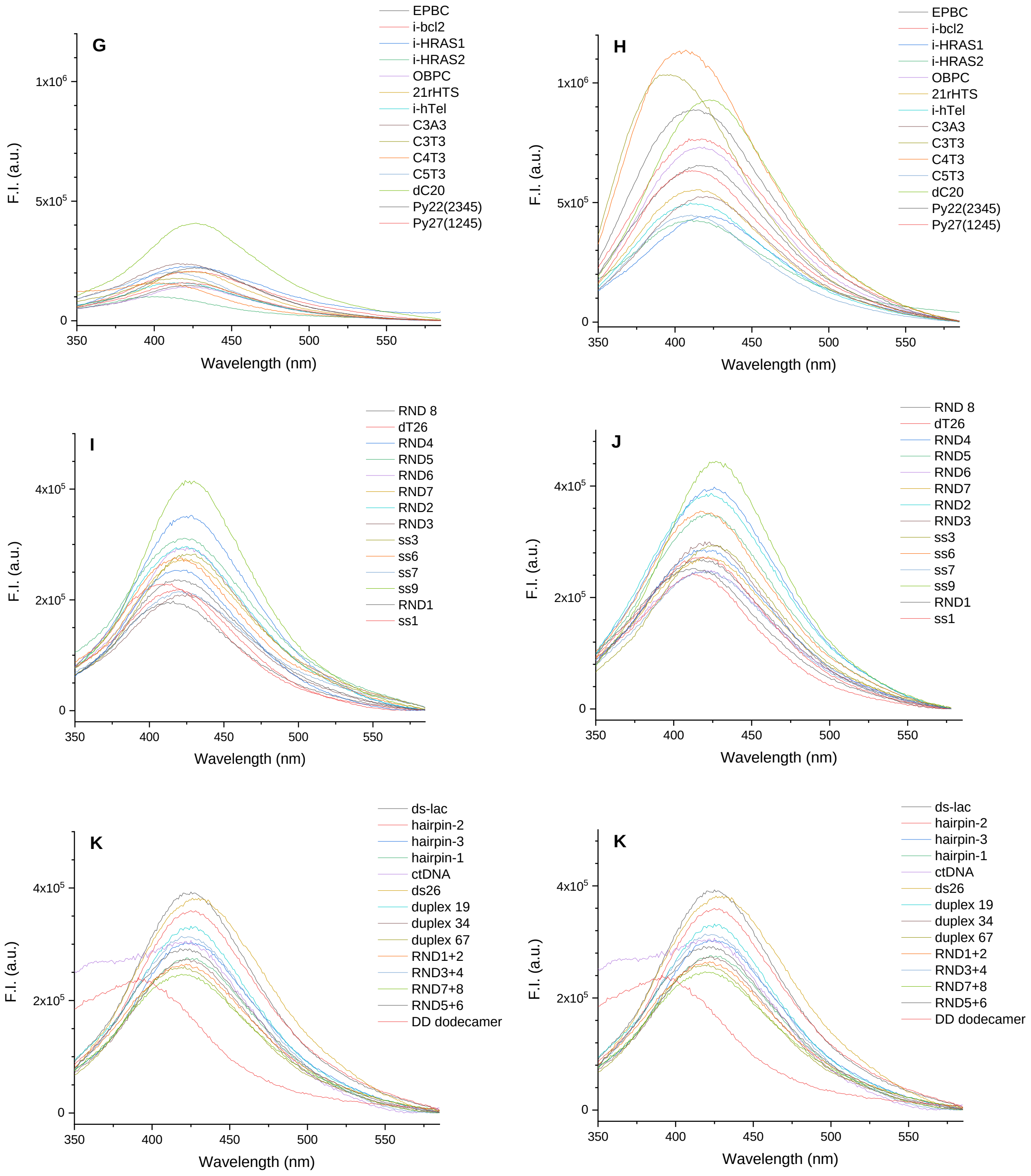


**Figure S9.** Emission spectra ($\lambda_{ex}$ = 300 nm) of the tested DNA sequences ($c$ = 5.7 µM), grouped according to their expected conformation: A–B) parallel G4; C–D) anti-parallel G4; E–F) hybrid G4; G–H) iM; I–J) single strands; K–L) duplex, in buffers A (A, C, E, G, I, K) and B (B, D, F, H, J, L).

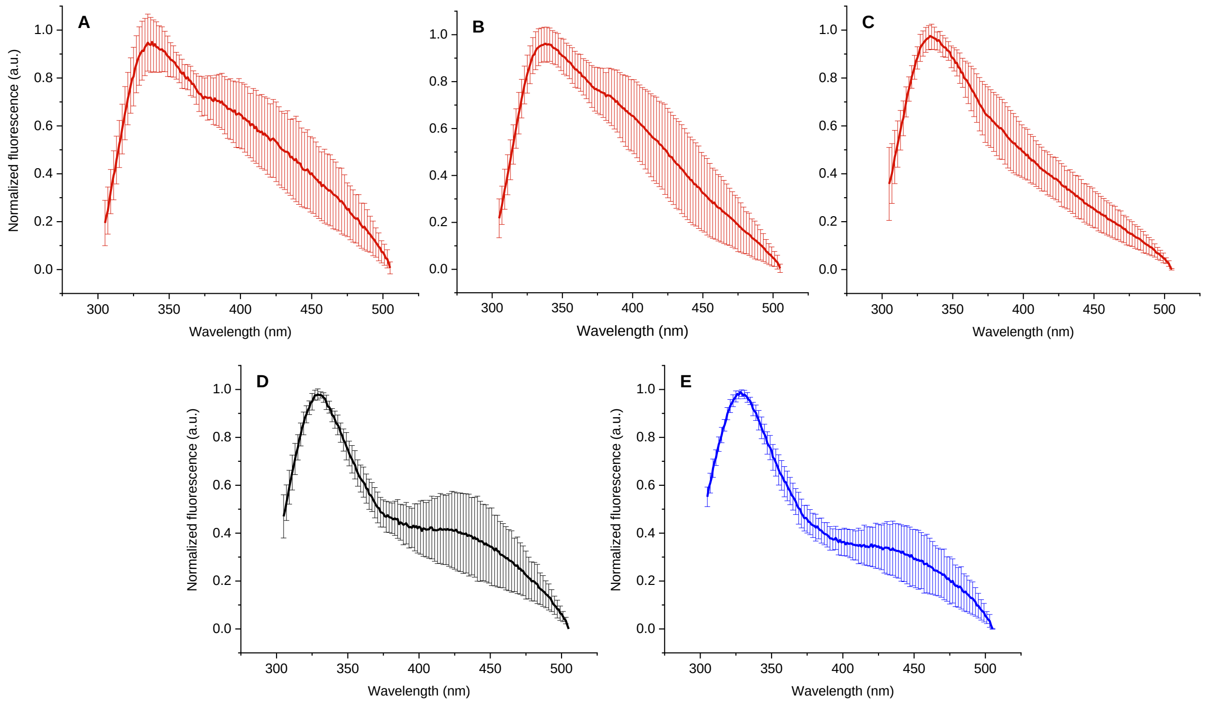


**Figure S10.** Group-averaged, normalized emission spectra ($\lambda_{ex}$ = 260 nm, buffer B) of the tested oligonucleotides, grouped according to their expected conformation: A) hybrid G4s; B) parallel G4s, C) anti-parallel G4s, D) single strands, E) duplexes. Error bars represent the standard deviation of the emission at a given wavelength inside each conformational group. *LWDLN-1*, *19wt* and *SP-PGQ3* were excluded from the analysis.

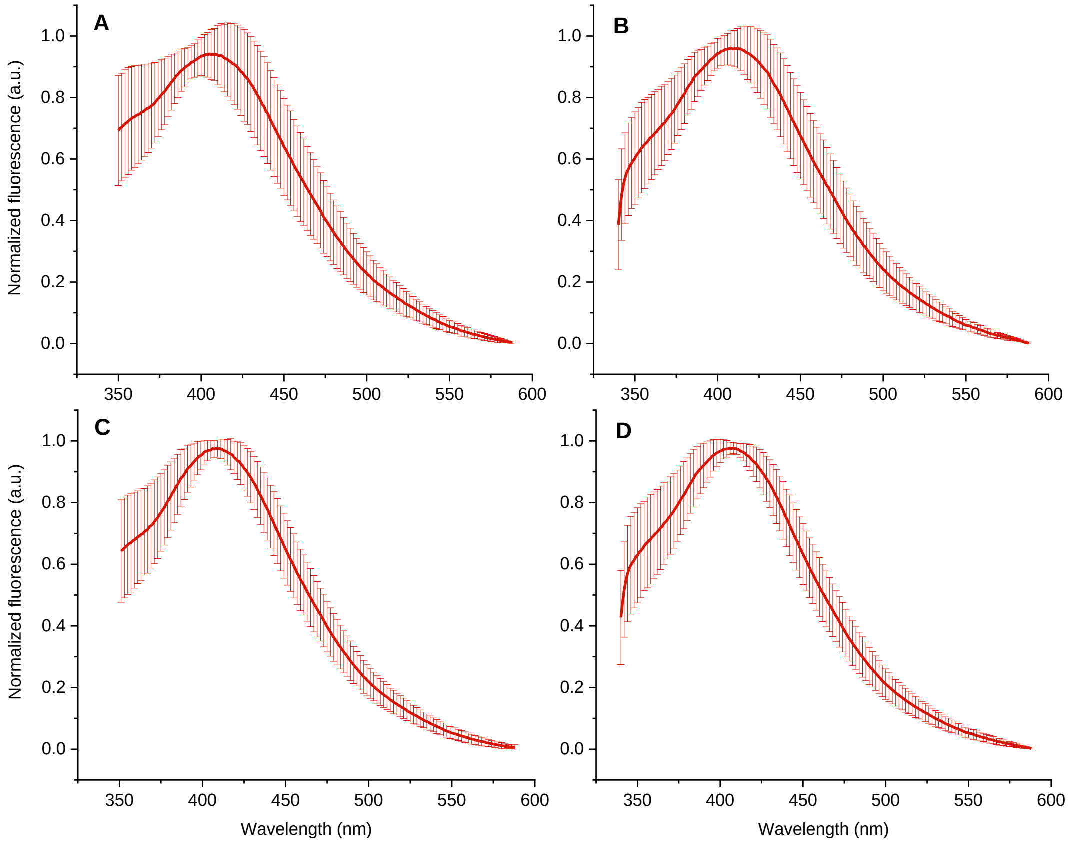

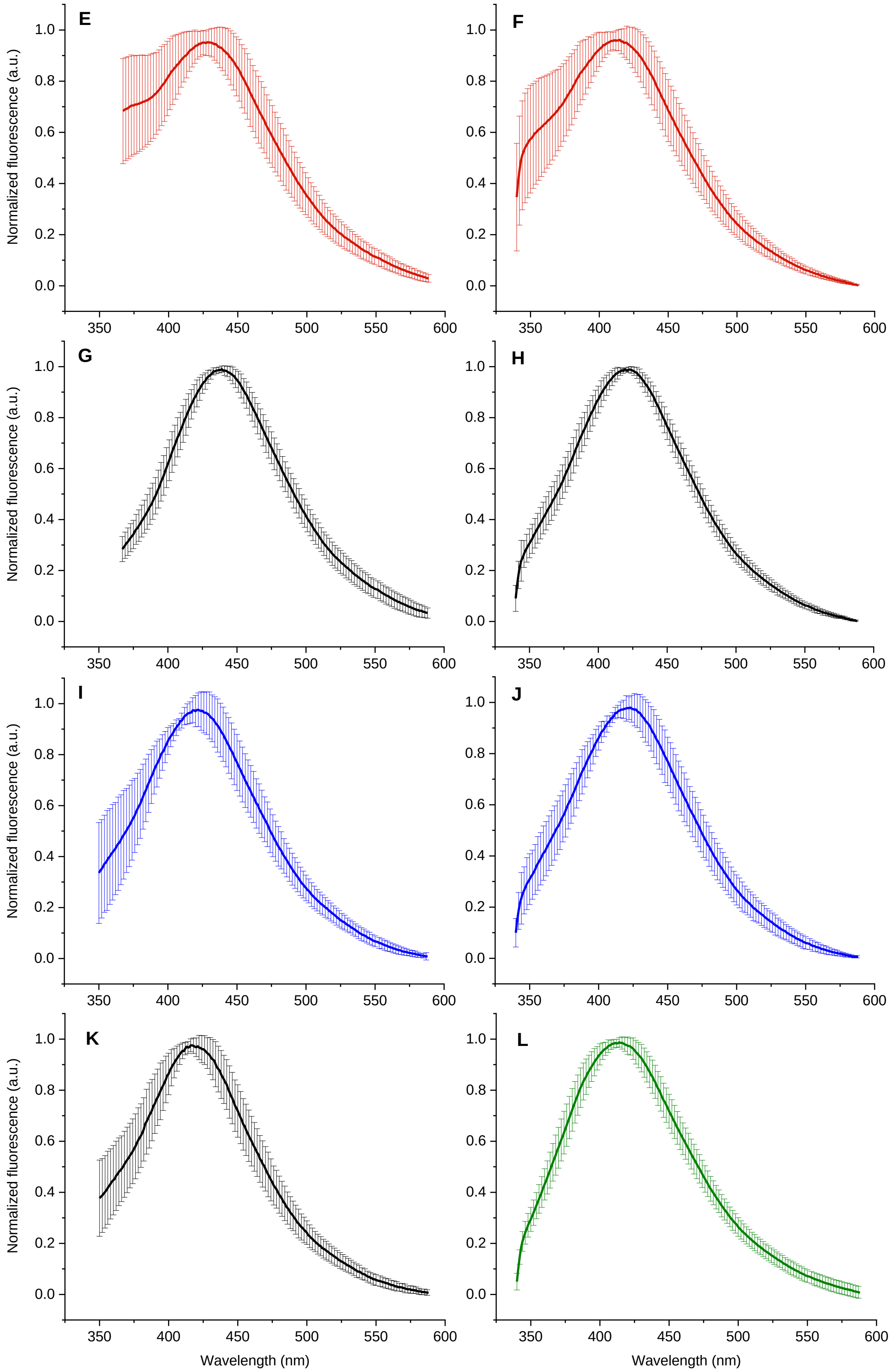


**Figure S11.** Group-averaged, normalized emission spectra ($\lambda_{ex}$ = 300 nm, buffers A and B) of the tested oligonucleotides, grouped according to their expected conformation. A–B) hybrid G4s; C–D) parallel G4s, E–F) anti-parallel G4s, G–H) single strands, I–J) duplexes, K–L) iM-forming sequences. Spectra presented in A, C, E, G, I, K were acquired in buffer A; spectra presented in B, D, F, H, J, L were acquired in buffer B. Error bars represent the standard deviation of the emission at a given wavelength inside each conformational group. *LWDLN-1*, *19wt* and *SP-PGQ3* were excluded from the analysis.

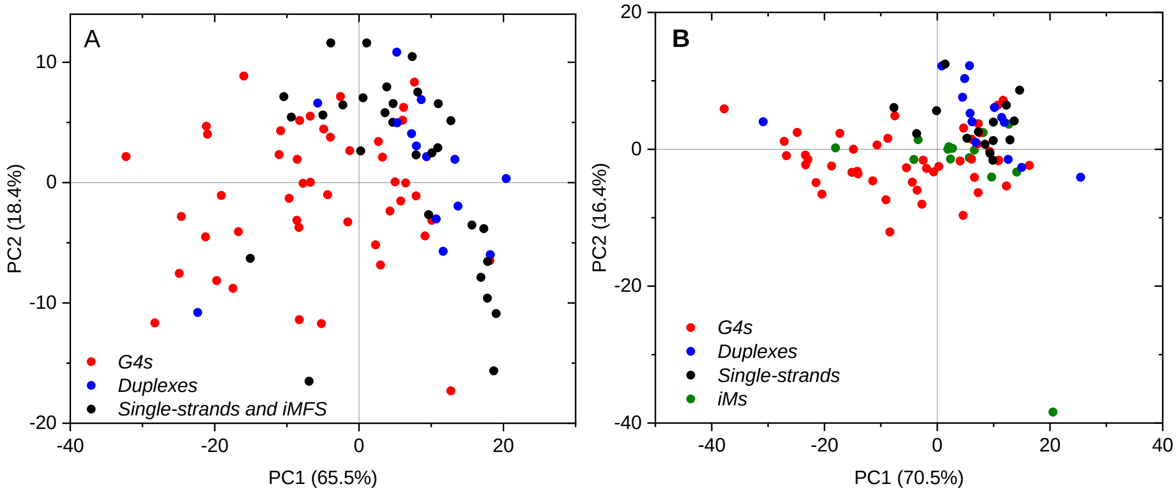


**Figure S12.** PC1 vs. PC2 plots resulting from the PCA of normalized emission spectra recorded A) in buffer A ($\lambda_{ex}$ = 300 nm); B) in buffer B ($\lambda_{ex}$ = 300 nm). In (A), the putative iM-forming sequences are shown are grouped with single-strands (in black), as they are mostly not folded in this buffer. In B, they are instead shown in green, since they adopt a distinct conformation.

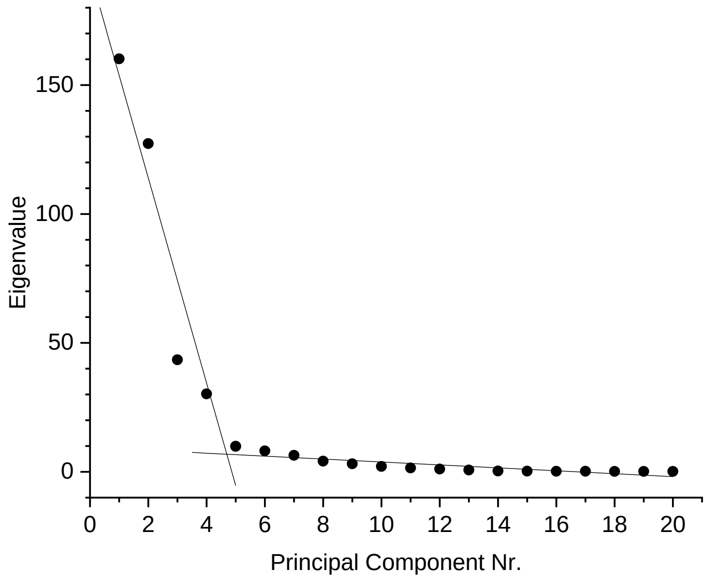


**Figure S13.** Scree plot of the eigenvalues obtained for the PCA shown in Figure 6, A of the main text. The change in slope observed at the level of PC5 indicates that the variance is mostly described by the first five components, whereas the contribution of the remaining 15 is negligible.

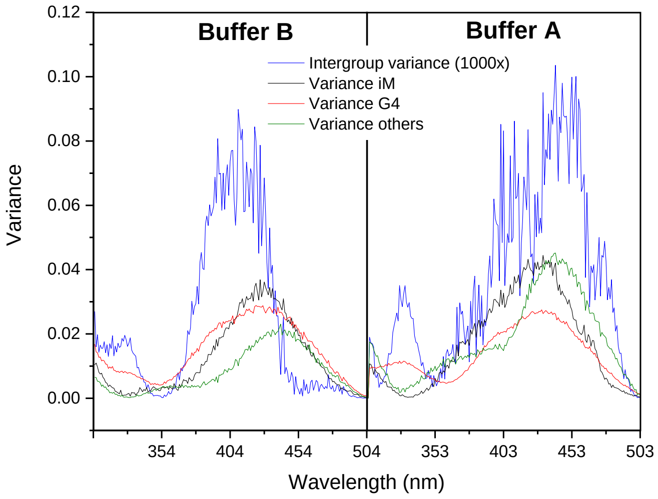


**Figure S14.** Analysis of the variance of the emission dataset. The variance within each group at every emission wavelength (305–505 nm in buffer A and buffer B, $\lambda_{ex}$ = 260 nm) is calculated from the normalized emission dataset and is plotted as a function of the emission wavelength. The inter-group variance is also calculated from the resulting three values and is plotted (1000x magnification) as a function of the emission wavelength.

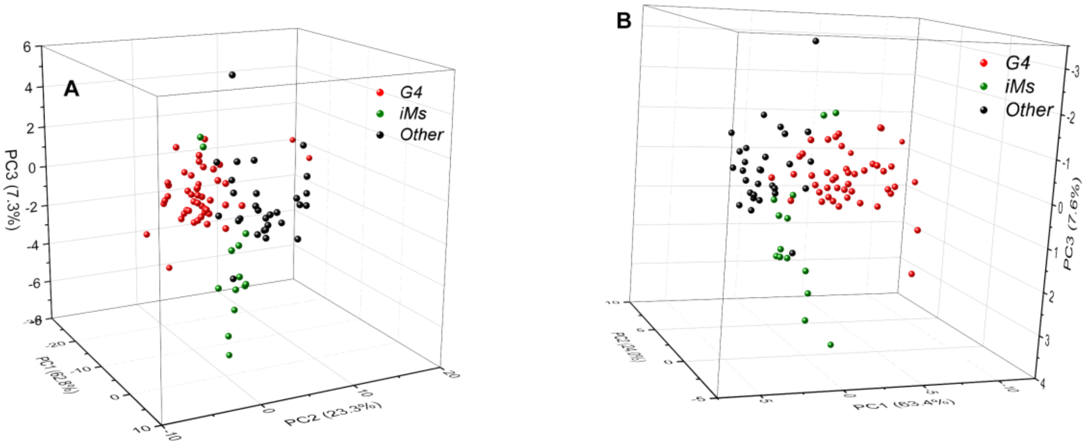


**Figure S15.** 3D PCA plot (PC1 vs. PC2 vs. PC3) resulting from A) analysis of the combination of normalized emission data recorded at 333–343 nm and 378–388 nm in buffer A and 324–333 nm and 374–393 nm in buffer B, upon excitation at 260 nm; B) analysis of the combination of normalized emission data recorded at 335, 338, 341, 380, 383 and 386 nm in buffer A and 329, 332, 376, 379, 382, 385, 388 and 391 nm in buffer B, upon excitation at 260 nm.

**Table S3.** Sequences and conformational assignment of test sequences *RND-HS1* to *RND-HS4*.

| Acronym | Sequence (5′–3′)[a] | G4 score [b] | Probability *P* [c] | | | Assign-ment [c] |
|---|---|---|---|---|---|---|
| | | | G4 | iM | Other | |
| *RND-HS1* | GAGTGAGGGGCTCGGCGCGGTGAGG | 1.16 | $5.5\times10^{-4}$ | $7.5\times10^{-6}$ | 0.99 | Other |
| *RND-HS2* | GC**CGCCCG**GGG**CGGGCG**CAGGGGGC | 1.24 | $9.5\times10^{-2}$ | $1.0\times10^{-4}$ | 0.90 | Other |
| *RND-HS3* | GGG**GCGCGC**GCA**GCGCGC**GGCGGG | 1.12 | 0.98 | $3.1\times10^{-3}$ | $1.3\times10^{-2}$ | G4 |
| *RND-HS4* | CCGGCTCGGGGATCGGGGGAGCGTC | 1.32 | 0.80 | $1.5\times10^{-3}$ | 0.19 | G4 |

[a] Consecutive runs of ≥ two guanines are underlined; self-complementary runs are printed in bold. [b] Calculated using the G4Hunter algorithm (https://bioinformatics.cruk.cam.ac.uk/G4Hunter/). [c] Based on the LDA analysis of the emission dataset.

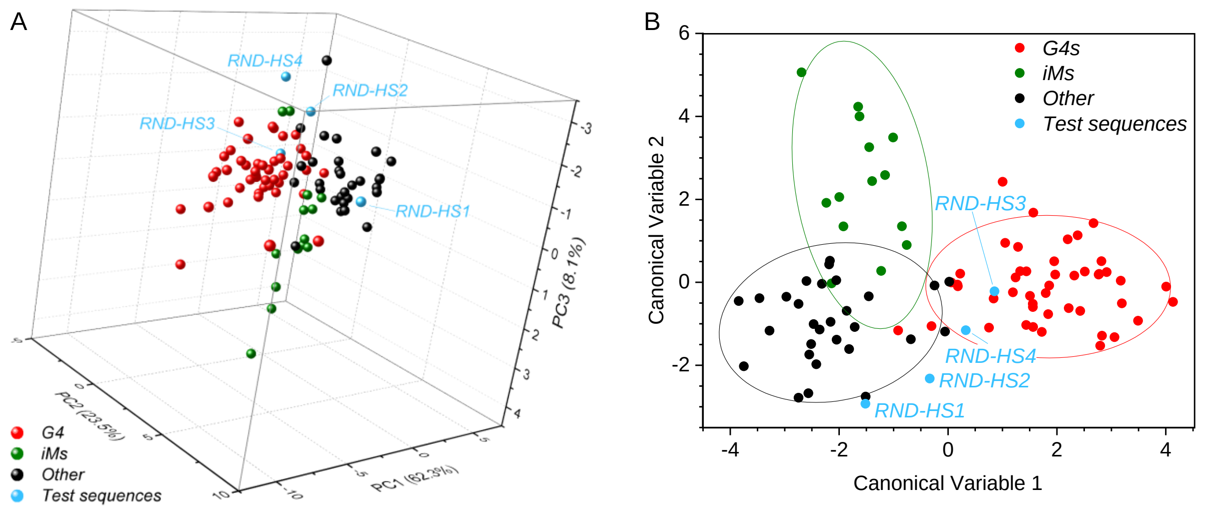


**Figure S16.** A) PC1 vs. PC2 vs. PC3 plot obtained from the analysis of the training set (reduced dataset), supplemented with the emission data obtained for test sequences *RND-HS1* to *4*. B) LDA plot for the same dataset.

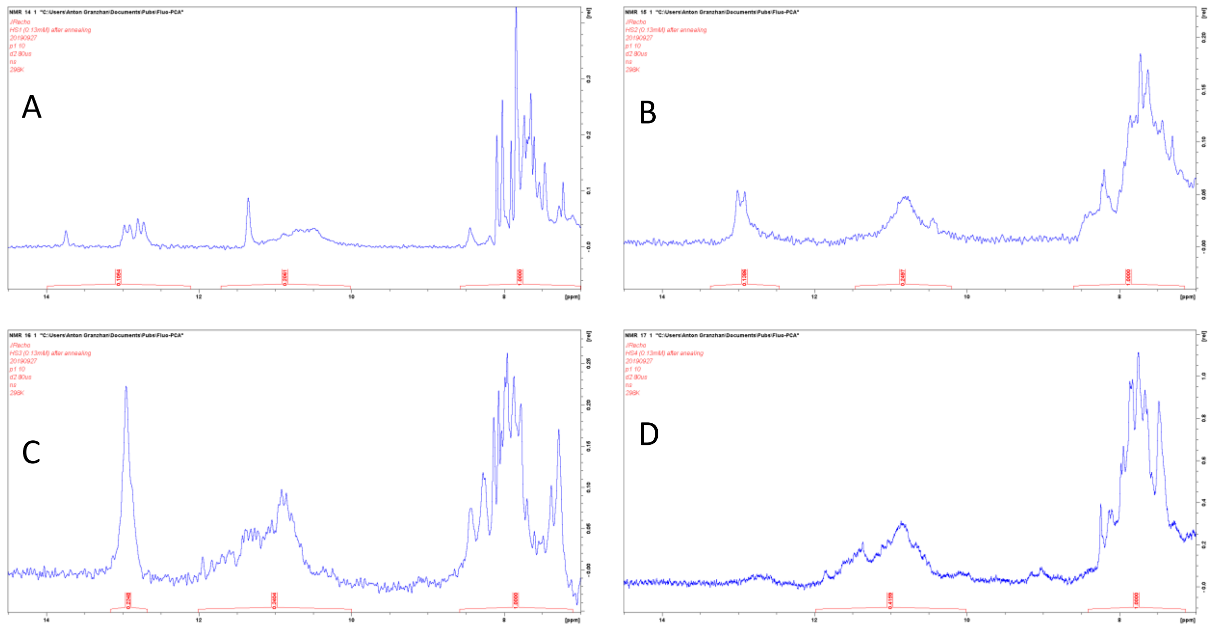


**Figure S17.** $^1$H NMR spectra (500 MHz) of oligonucleotides *RND-HS1* (A), *RND-HS2* (B), *RND-HS3* (C), and *RND-HS4* (D) in Buffer A (containing 20% v/v $D_2O$); *c* = 130 μM in all cases.

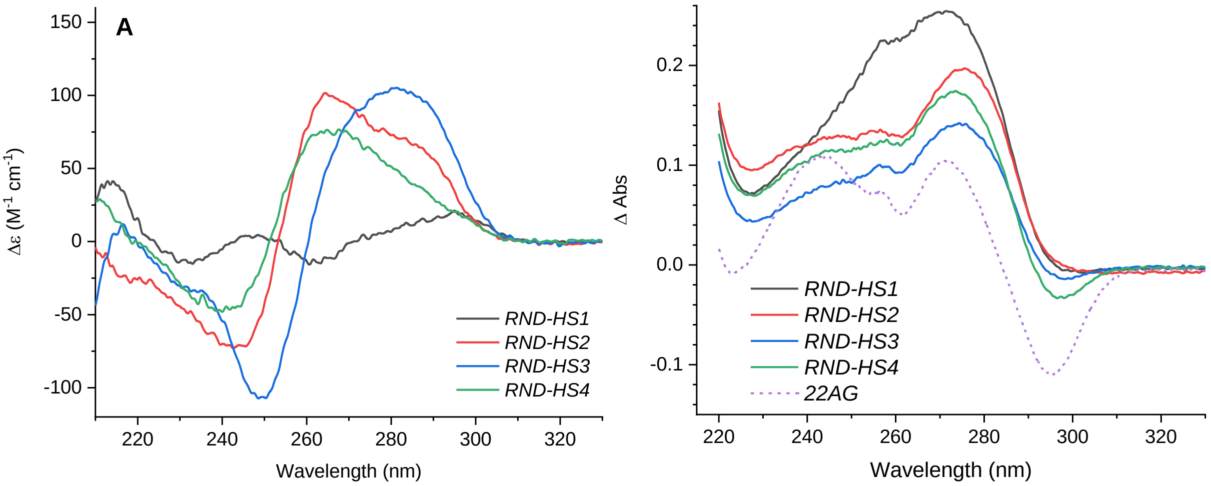


**Figure S18.** A) CD and B) thermal difference spectra (TDS, ΔAbs = $Abs_{bs90\ °C}$ – $Abs_{20\ °C}$) of test sequences *RND-HS1* to *RND-HS4* in buffer A ($c$ = 5.7 µM). A TDS of a typical G4 structure (*22AG*) is included for comparison in B).

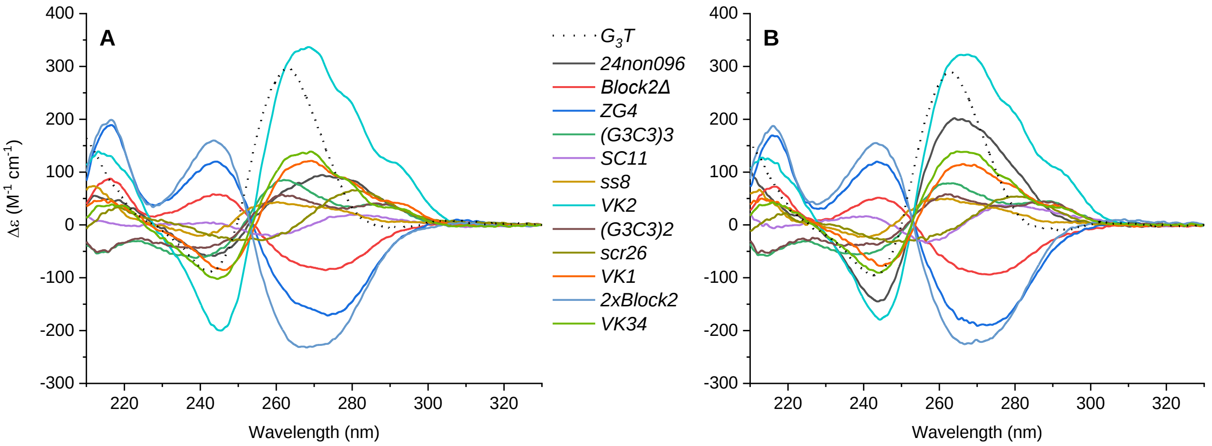


**Figure S19.** CD spectra of oligonucleotides adopting peculiar conformations A) in buffer A and B) in buffer B ($c$ = 5.7 µM).

**Table S4.** LDA-based assignment of the test sequences presented in Figure S18.

| Acronym | Sequence (5′–3′) | Structure (PDB) | Probability *P* | | | Assign-ment |
|---|---|---|---|---|---|---|
| | | | **G4** | **i-motif** | **Other** | |
| *G₃T* | GGGTGGGTGGGTGGG | stacked dimer G4 (**2LE6**)[a] | **1** | $6.8\times10^{-10}$ | $7.0\times10^{-6}$ | G4 |
| *ZG4* | TGGTGGTGGTGGTTGTGGTGGTG GTGTT | Z-G4 (**2MS9**) | **1** | $1.9\times10^{-12}$ | $3.5\times10^{-7}$ | G4 |
| *Block2Δ* | GTGGTGGTGGTG | Z-G4 (**6FQ2**) | **1** | $2.3\times10^{-11}$ | $4.8\times10^{-7}$ | G4 |
| *2xBlock2* | GTGGTGGTGGTGTTGTGGTGGTG GTGT | Z-G4 (**6GZ6**) | **1** | $1.7\times10^{-12}$ | $5.7\times10^{-7}$ | G4 |
| *VK1* | GGGAGCGAGGGAGCG | non-G4 quad-ruplex (**2MJJ**) | **0.98** | $2.1\times10^{-3}$ | $2.1\times10^{-2}$ | G4 |
| *VK2* | GGGAGCGAGGGAGCGAGGGAGC GAGGGAGCG | non-G4 quad-ruplex | **0.98** | $1.4\times10^{-2}$ | $7.9\times10^{-3}$ | G4 |
| *VK34* | GCGAGGGAGCGAGGG | non-G4 quad-ruplex (**5M1L**) | $3.8\times10^{-3}$ | $6.4\times10^{-2}$ | **0.93** | Other |
| *$(G_3C_3)_3$* | TGGGCCCGGGCCCGGGCCC | A-type duplex | $5.7\times10^{-5}$ | $1.3\times10^{-3}$ | **0.99** | Other |
| *$(G_3C_3)_2$* | TGGGCCCGGGCCC | A-type duplex | $4.0\times10^{-6}$ | $1.9\times10^{-4}$ | **0.99** | Other |
| *SC11* | GTGTGGGTGTG | G-hairpin (**5M1W**) | $2.2\times10^{-2}$ | $3.1\times10^{-3}$ | **0.97** | Other |
| *ss8* | GGAGAGAGAGTGTGTGTGTGGG | unknown | $6.2\times10^{-7}$ | **0.87** | **0.13** | iM |
| *24non096* | TGGGATGCGACAGAGAGGACGG GA | unknown | $3.7\times10^{-2}$ | $2.5\times10^{-2}$ | **0.94** | Other |
| *scr26* | GAAGTGTGTGTGTGTGTGTGTGT GAA | unknown | $4.2\times10^{-4}$ | **0.48** | **0.51** | Other |

[a] PDB entry for the inosine-modified analogue (*J19*, 5′-GIGTGGGTGGGTGGGT-3′).

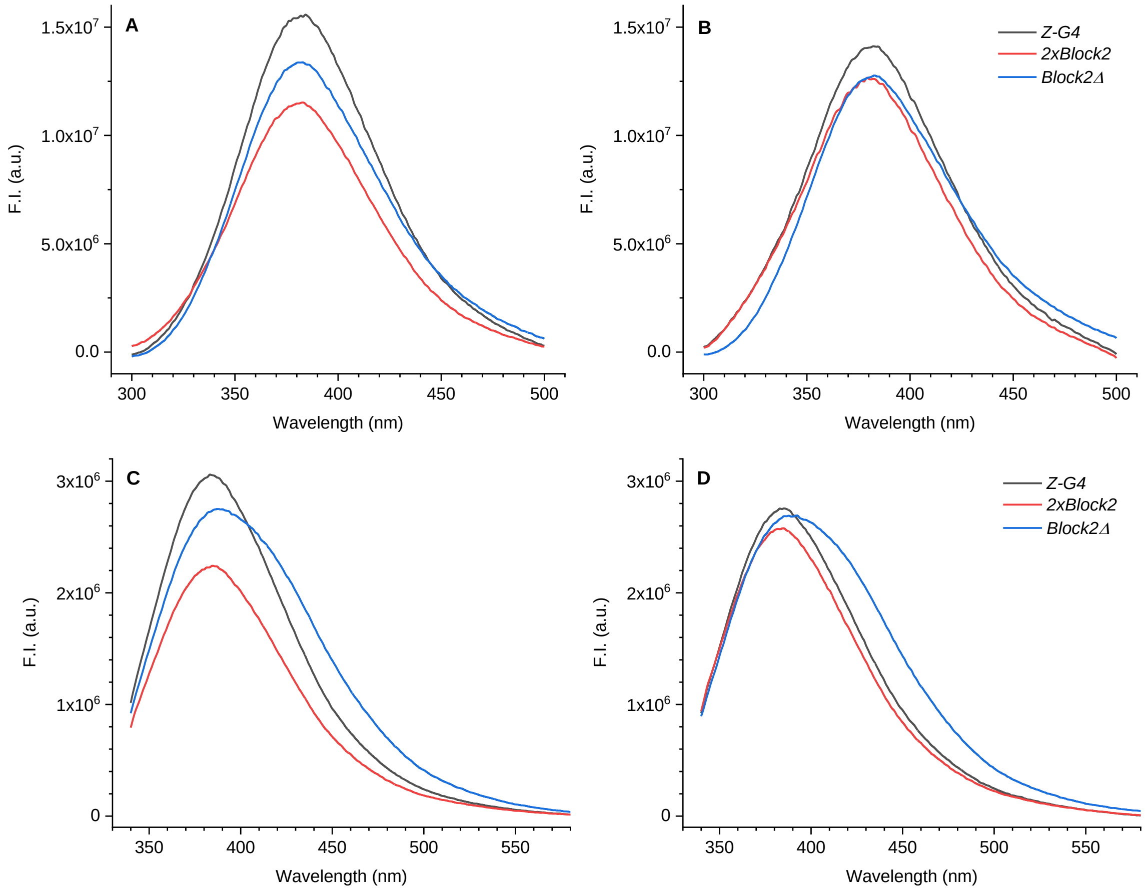


**Figure S20.** Fluorescence emission spectra of left-handed G4s (*ZG4* and *2xBlock2*: $c$ = 5.7 µM, *Block2Δ*: $c$ = 11.4 µM), A) in buffer A, $\lambda_{ex}$ = 260 nm; B) in buffer B, $\lambda_{ex}$ = 260 nm; C) in buffer A, $\lambda_{ex}$ = 300 nm; B) in buffer B, $\lambda_{ex}$ = 300 nm. Spectra were corrected for the inner-filter effect.

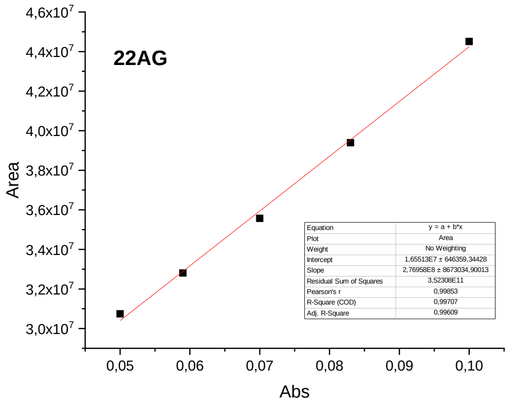
22AG
Area
Abs
$4,6x10^7$
$4,4x10^7$
$4,2x10^7$
$4,0x10^7$
$3,8x10^7$
$3,6x10^7$
$3,4x10^7$
$3,2x10^7$
$3,0x10^7$
0,05
0,06
0,07
0,08
0,09
0,10
Equation y = a + b*x
Plot Area
Weight No Weighting
Intercept 1,65513E7 ± 646359,34428
Slope 2,76958E8 ± 8673034,90013
Residual Sum of Squares 3,52308E11
Pearson's r 0,99853
R-Square (COD) 0,99707
Adj. R-Square 0,99609

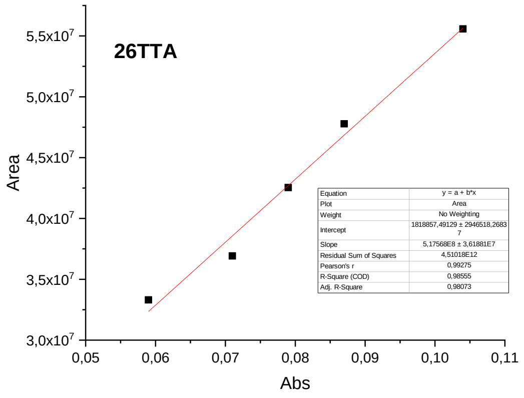
26TTA
Area
Abs
$5,5x10^7$
$5,0x10^7$
$4,5x10^7$
$4,0x10^7$
$3,5x10^7$
$3,0x10^7$
0,05
0,06
0,07
0,08
0,09
0,10
0,11
Equation y = a + b*x
Plot Area
Weight No Weighting
Intercept 1818857,49129 ± 2946518,26837
Slope 5,17568E8 ± 3,61881E7
Residual Sum of Squares 4,51018E12
Pearson's r 0,99275
R-Square (COD) 0,98555
Adj. R-Square 0,98073

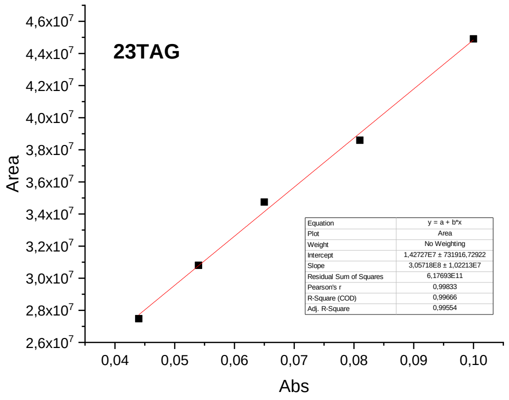
23TAG
Area
Abs
$4,6x10^7$
$4,4x10^7$
$4,2x10^7$
$4,0x10^7$
$3,8x10^7$
$3,6x10^7$
$3,4x10^7$
$3,2x10^7$
$3,0x10^7$
$2,8x10^7$
$2,6x10^7$
0,04
0,05
0,06
0,07
0,08
0,09
0,10
Equation y = a + b*x
Plot Area
Weight No Weighting
Intercept 1,42727E7 ± 731916,72922
Slope 3,05718E8 ± 1,02213E7
Residual Sum of Squares 6,17693E11
Pearson's r 0,99833
R-Square (COD) 0,99666
Adj. R-Square 0,99554

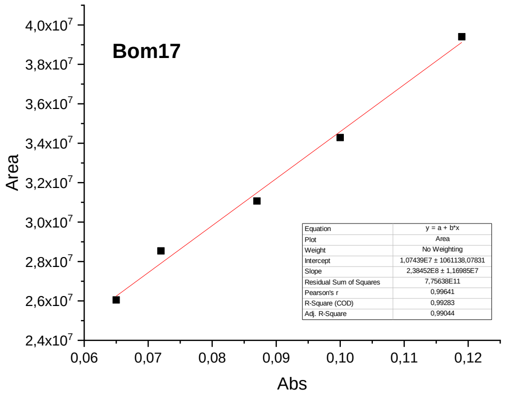
Bom17
Area
Abs
$4,0x10^7$
$3,8x10^7$
$3,6x10^7$
$3,4x10^7$
$3,2x10^7$
$3,0x10^7$
$2,8x10^7$
$2,6x10^7$
$2,4x10^7$
0,06
0,07
0,08
0,09
0,10
0,11
0,12
Equation y = a + b*x
Plot Area
Weight No Weighting
Intercept 1,07439E7 ± 1061138,07831
Slope 2,38452E8 ± 1,16985E7
Residual Sum of Squares 7,75638E11
Pearson's r 0,99641
R-Square (COD) 0,99283
Adj. R-Square 0,99044

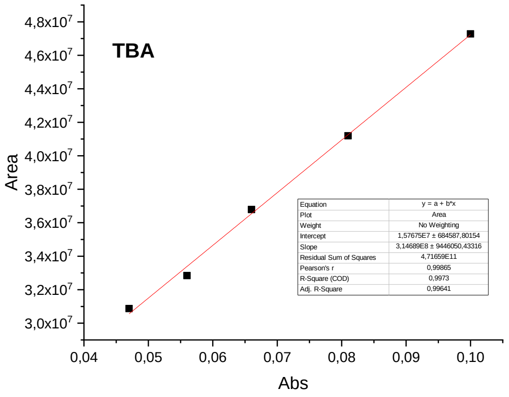
TBA
Area
Abs
$4,8x10^7$
$4,6x10^7$
$4,4x10^7$
$4,2x10^7$
$4,0x10^7$
$3,8x10^7$
$3,6x10^7$
$3,4x10^7$
$3,2x10^7$
$3,0x10^7$
0,04
0,05
0,06
0,07
0,08
0,09
0,10
Equation y = a + b*x
Plot Area
Weight No Weighting
Intercept 1,57675E7 ± 684587,80154
Slope 3,14689E8 ± 9446050,43316
Residual Sum of Squares 4,71659E11
Pearson's r 0,99865
R-Square (COD) 0,9973
Adj. R-Square 0,99641

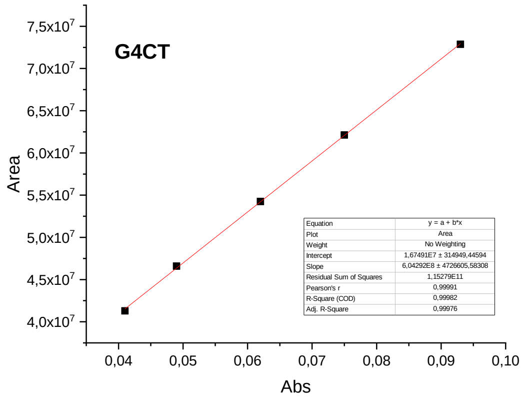
G4CT
Area
Abs
$7,5x10^7$
$7,0x10^7$
$6,5x10^7$
$6,0x10^7$
$5,5x10^7$
$5,0x10^7$
$4,5x10^7$
$4,0x10^7$
0,04
0,05
0,06
0,07
0,08
0,09
0,10
Equation y = a + b*x
Plot Area
Weight No Weighting
Intercept 1,67491E7 ± 314949,44594
Slope 6,04292E8 ± 4726605,58308
Residual Sum of Squares 1,15279E11
Pearson's r 0,99991
R-Square (COD) 0,99982
Adj. R-Square 0,99976

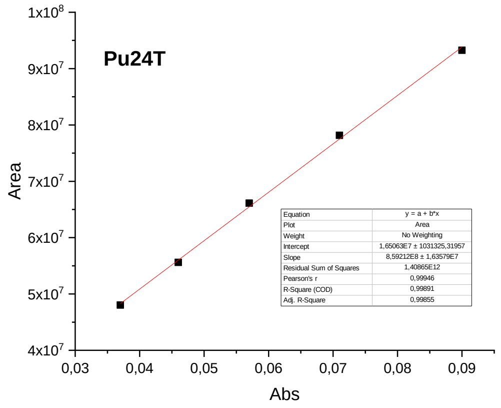
Pu24T
Area
Abs
$1x10^8$
$9x10^7$
$8x10^7$
$7x10^7$
$6x10^7$
$5x10^7$
$4x10^7$
0,03
0,04
0,05
0,06
0,07
0,08
0,09
Equation y = a + b*x
Plot Area
Weight No Weighting
Intercept 1,65063E7 ± 1031325,31957
Slope 8,59212E8 ± 1,63579E7
Residual Sum of Squares 1,40865E12
Pearson's r 0,99946
R-Square (COD) 0,99891
Adj. R-Square 0,99855

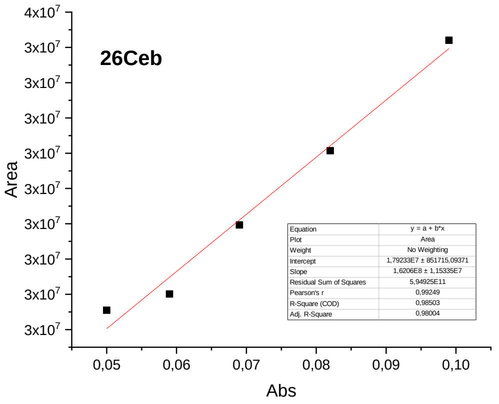
26Ceb
Area
Abs
$4x10^7$
$3x10^7$
$3x10^7$
$3x10^7$
$3x10^7$
$3x10^7$
$3x10^7$
$3x10^7$
$3x10^7$
$3x10^7$
0,05
0,06
0,07
0,08
0,09
0,10
Equation y = a + b*x
Plot Area
Weight No Weighting
Intercept 1,79233E7 ± 851715,09371
Slope 1,6206E8 ± 1,15335E7
Residual Sum of Squares 5,94925E11
Pearson's r 0,99249
R-Square (COD) 0,98503
Adj. R-Square 0,98004

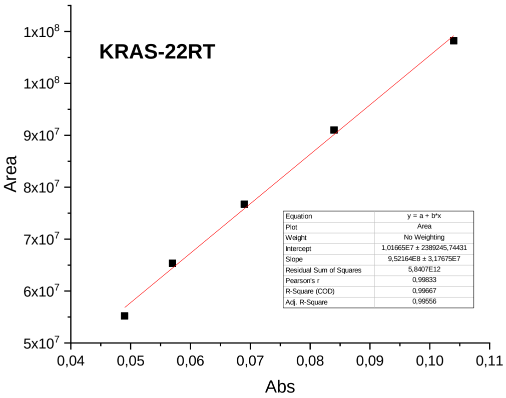
KRAS-22RT
Area
Abs
1x10^8
1x10^8
9x10^7
8x10^7
7x10^7
6x10^7
5x10^7
0,04 0,05 0,06 0,07 0,08 0,09 0,10 0,11
Equation y = a + b*x
Plot Area
Weight No Weighting
Intercept 1,01665E7 ± 2389245,74431
Slope 9,52164E8 ± 3,17675E7
Residual Sum of Squares 5,8407E12
Pearson's r 0,99833
R-Square (COD) 0,99667
Adj. R-Square 0,99556

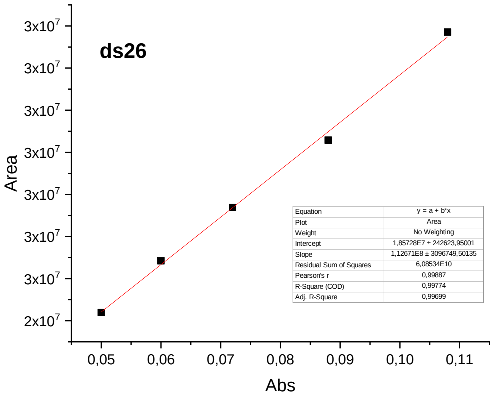
ds26
Area
Abs
3x10^7
3x10^7
3x10^7
3x10^7
3x10^7
3x10^7
3x10^7
2x10^7
0,05 0,06 0,07 0,08 0,09 0,10 0,11
Equation y = a + b*x
Plot Area
Weight No Weighting
Intercept 1,85728E7 ± 242623,95001
Slope 1,12671E8 ± 3096749,50135
Residual Sum of Squares 6,08534E10
Pearson's r 0,99887
R-Square (COD) 0,99774
Adj. R-Square 0,99699

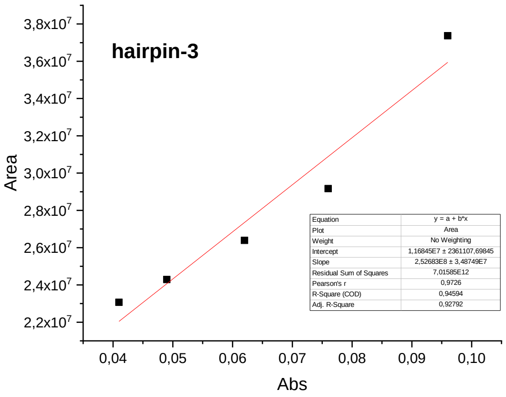
hairpin-3
Area
Abs
3,8x10^7
3,6x10^7
3,4x10^7
3,2x10^7
3,0x10^7
2,8x10^7
2,6x10^7
2,4x10^7
2,2x10^7
0,04 0,05 0,06 0,07 0,08 0,09 0,10
Equation y = a + b*x
Plot Area
Weight No Weighting
Intercept 1,16845E7 ± 2361107,69845
Slope 2,52683E8 ± 3,48749E7
Residual Sum of Squares 7,01585E12
Pearson's r 0,9726
R-Square (COD) 0,94594
Adj. R-Square 0,92792

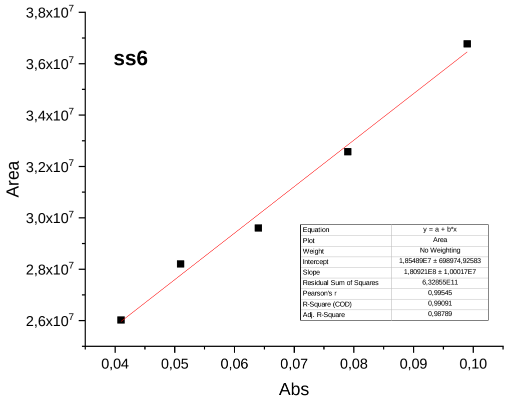
ss6
Area
Abs
3,8x10^7
3,6x10^7
3,4x10^7
3,2x10^7
3,0x10^7
2,8x10^7
2,6x10^7
0,04 0,05 0,06 0,07 0,08 0,09 0,10
Equation y = a + b*x
Plot Area
Weight No Weighting
Intercept 1,85489E7 ± 698974,92583
Slope 1,80921E8 ± 1,00017E7
Residual Sum of Squares 6,32855E11
Pearson's r 0,99545
R-Square (COD) 0,99091
Adj. R-Square 0,98789

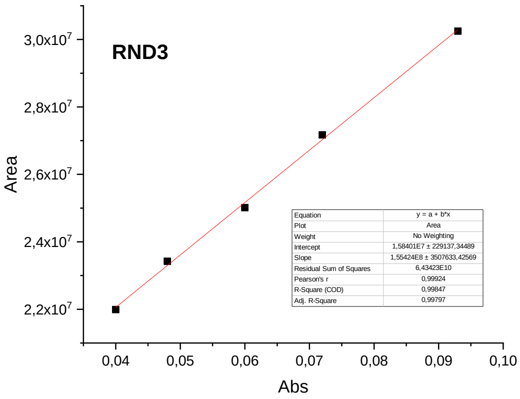
RND3
Area
Abs
3,0x10^7
2,8x10^7
2,6x10^7
2,4x10^7
2,2x10^7
0,04 0,05 0,06 0,07 0,08 0,09 0,10
Equation y = a + b*x
Plot Area
Weight No Weighting
Intercept 1,58401E7 ± 229137,34489
Slope 1,55424E8 ± 3507633,42569
Residual Sum of Squares 6,43423E10
Pearson's r 0,99924
R-Square (COD) 0,99847
Adj. R-Square 0,99797

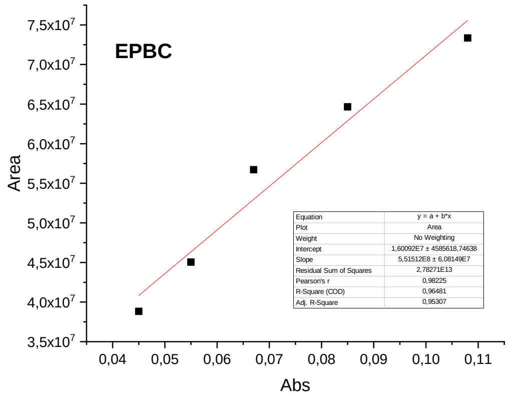
EPBC
Area
Abs
7,5x10^7
7,0x10^7
6,5x10^7
6,0x10^7
5,5x10^7
5,0x10^7
4,5x10^7
4,0x10^7
3,5x10^7
0,04 0,05 0,06 0,07 0,08 0,09 0,10 0,11
Equation y = a + b*x
Plot Area
Weight No Weighting
Intercept 1,60092E7 ± 4585618,74638
Slope 5,51512E8 ± 6,08149E7
Residual Sum of Squares 2,78271E13
Pearson's r 0,98225
R-Square (COD) 0,96481
Adj. R-Square 0,95307

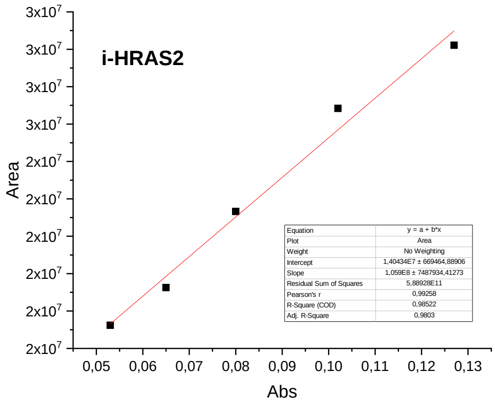
i-HRAS2
Area
Abs
3x10^7
3x10^7
3x10^7
3x10^7
2x10^7
2x10^7
2x10^7
2x10^7
2x10^7
2x10^7
0,05 0,06 0,07 0,08 0,09 0,10 0,11 0,12 0,13
Equation y = a + b*x
Plot Area
Weight No Weighting
Intercept 1,40434E7 ± 669464,88906
Slope 1,059E8 ± 7487934,41273
Residual Sum of Squares 5,88928E11
Pearson's r 0,99258
R-Square (COD) 0,98522
Adj. R-Square 0,9803

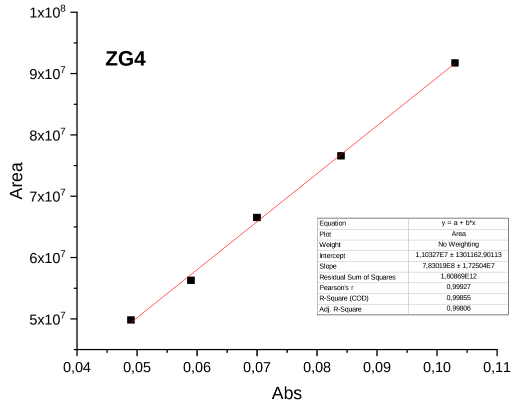
ZG4
Area
Abs
1x10^8
9x10^7
8x10^7
7x10^7
6x10^7
5x10^7
0,04 0,05 0,06 0,07 0,08 0,09 0,10 0,11
Equation y = a + b*x
Plot Area
Weight No Weighting
Intercept 1,10327E7 ± 1301162,90113
Slope 7,83019E8 ± 1,72504E7
Residual Sum of Squares 1,60869E12
Pearson's r 0,99927
R-Square (COD) 0,99855
Adj. R-Square 0,99806

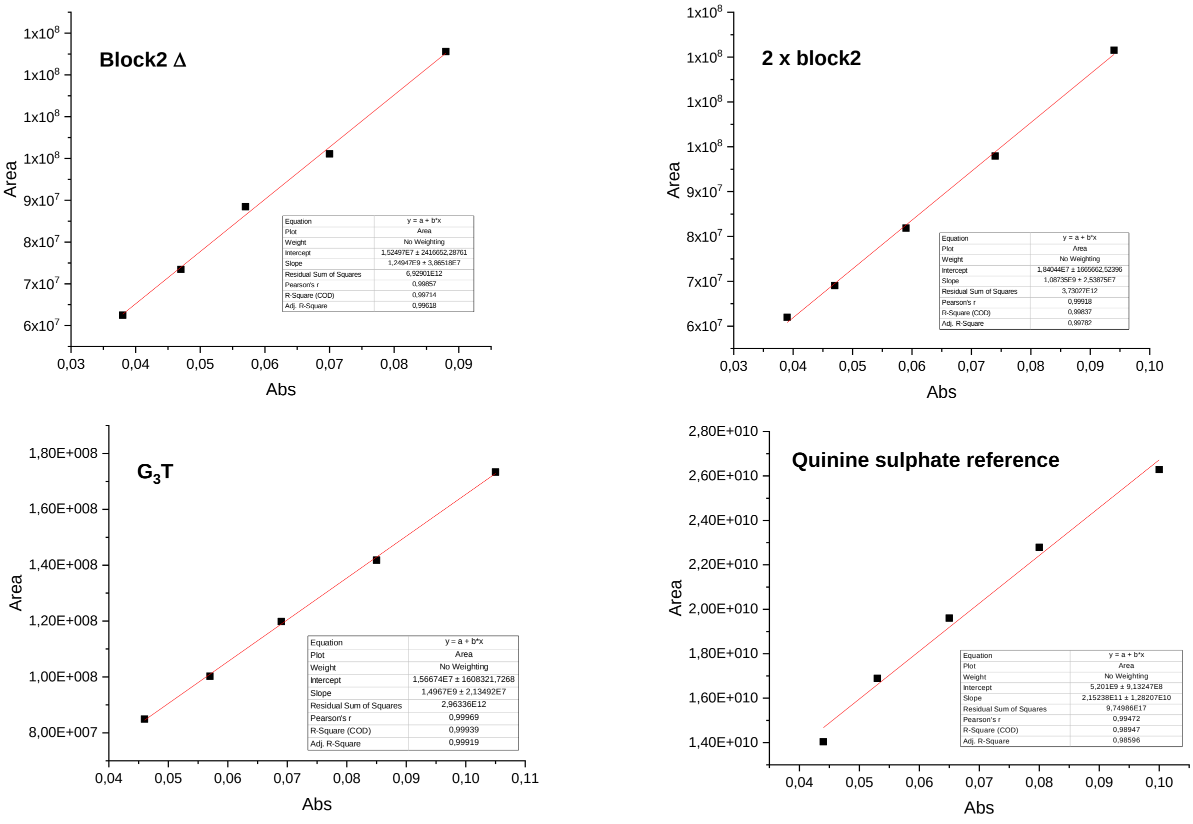


**Figure S21.** Determination of fluorescence quantum yields: plots of integrated fluorescence intensity vs. absorbance (at 265 nm) for serial dilutions of solutions of selected oligonucleotides. All experiments were performed in buffer A, except for EPBC and i-HRAS2 (buffer B), with initial solutions having $A \leq 0.11$ at the excitation wavelength. Data on quinine sulfate (reference material, $\Phi = 0.546$) were obtained in 0.5 M sulfuric acid.

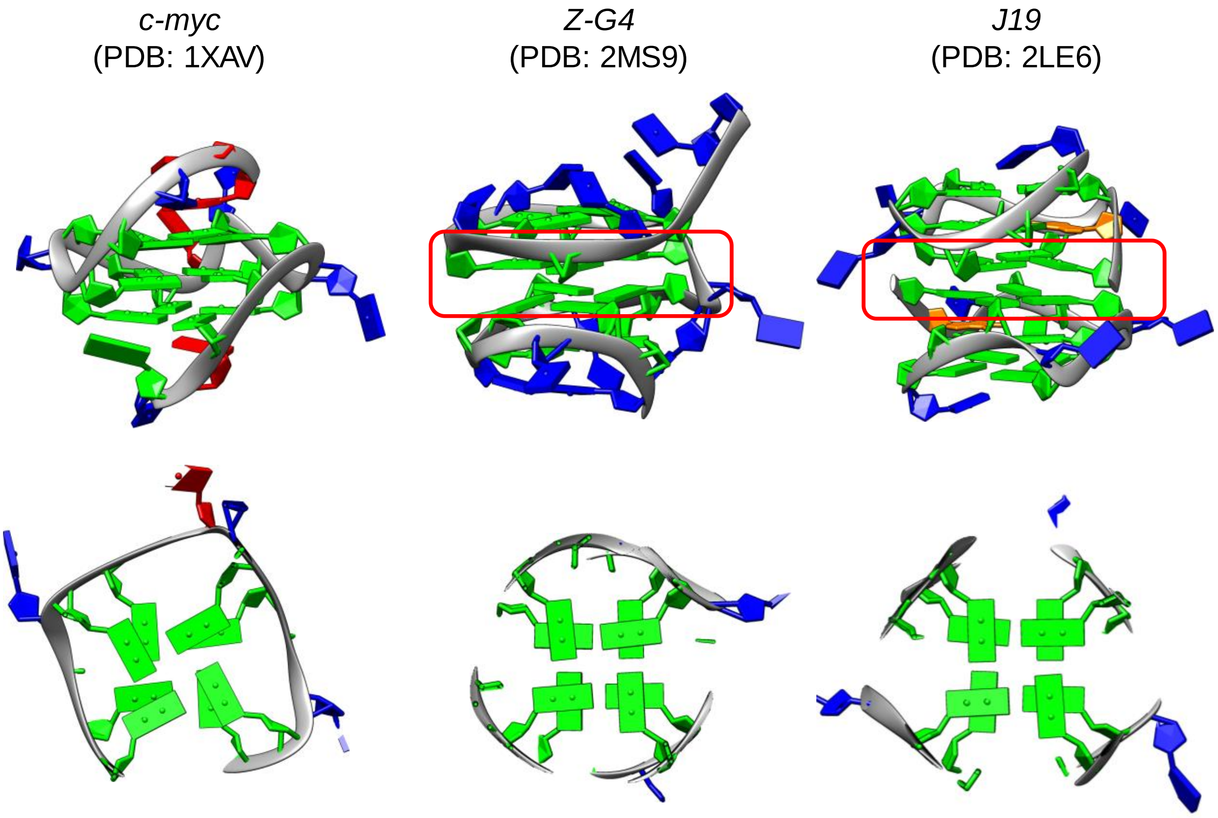


**Figure S22.** Lateral view of the whole G4 structure (top) and top view of G-quartet stacks (bottom) for a typical parallel G4 (*c-myc*), left-handed G4 (*ZG4*) and a stacked parallel dimeric G4 (*J19*, an inosine-modified analogue of $G_3T$). All three quadruplexes present only guanines in an *anti* orientation. However, both *ZG4* and *J19* present a characteristic 5/6-ring stacking mode of G-tetrads at the 5′–5′ interface of G4 blocks (highlighted in red). Nucleotide colors: dG, green; dA, red; dT, blue; dI, orange. Images generated with UCSF Chimera (*developed by the Resource for Biocomputing, Visualization, and Informatics at the University of California, San Francisco, with support from NIH P41-GM103311*).
